# Glutamate signaling at cytoneme synapses

**DOI:** 10.1101/475665

**Authors:** Hai Huang, Songmei Liu, Thomas B. Kornberg

**Affiliations:** Cardiovascular Research Institute, University of California San Francisco, San Francisco, CA2 94143

## Abstract

We investigated the roles of neuronal synapse components for development of the Drosophila air sac primordium (ASP). The ASP, an epithelial tube, extends specialized signaling filopodia called cytonemes that take up signals such as Dpp from the wing imaginal disc. Dpp signaling in the ASP requires that disc cells express Dpp, Synaptobrevin, Synaptotagmin-1, the glutamate transporter, and a voltage-gated calcium channel, and that ASP cells express the Dpp receptor, Synaptotagmin-4 and the AMPA-type glutamate receptor GluRII. Calcium transients in ASP cytonemes correlate with signaling activity. Calcium transients in the ASP require GluRII, are activated by L-glutamate and by stimulation of an optogenetic ion channel expressed in the wing disc, and are inhibited by EGTA and NASPM. Activation of GluRII is essential but not sufficient for signaling. Cytoneme-mediated signaling is glutamatergic.

**Summary:** Paracrine signals transfer between Drosophila epithelial cells at glutamatergic synapses.

## Introduction

Metazoan tissues are patterned by signaling proteins such as Bone morphogenetic protein (BMP), Fibroblast growth factor (FGF), Epidermal growth factor (EGF), Wnt, and Hedgehog. Our studies of the Drosophila air sac primordium (ASP), a single cell layered epithelial tube of the larval tracheal system, show that although it is patterned in part by Decapentaplegic (Dpp, a BMP) and Branchless (Bnl, a FGF), it does not produce these proteins (1, 2). The ASP does express the respective receptors for Dpp and Bnl/FGF, Thickveins (Tkv) and Breathless (Btl), and it receives Dpp and Bnl/FGF produced and secreted by the wing imaginal disc. The ASP develops adjacent to the wing disc, receiving Dpp and Bnl/FGF at distances of 5-40 μm. Despite the physical separation between disc cells that release these proteins and ASP cells that take them up, the proteins transfer to the ASP at cell-cell contacts. Specialized filopodia called cytonemes extend from ASP cells and synapse with disc cells (2, 3).

In addition to the ASP, cytoneme-mediated signaling has been implicated in Drosophila tissues such as the wing disc epithelium (4–8), abdominal histoblasts (6), testis (9), and the hematopoietic stem cell niche (10), and in zebrafish embryos (11–13), chick embryos (14), mouse embryos (15), and cultured cell systems (16). In signal producing cells, cytoneme-mediated signaling is dependent on the signaling proteins and requires proteins that process and prepare the signaling proteins for release (17). In cells that are situated between cells that produce signals and cells that extend cytonemes to receive them, components of the extracellular matrix and components of the planar polarity system are required (6, 18). And in receiving cells, signaling protein receptors and cytoskeletal, cell adhesion, and vesicle processing proteins are required (2, 5–7, 17, 18). In the ASP, mutants deficient for these proteins do not have cytonemes that extend to contact the disc, do not take up the signaling proteins, do not activate Dpp and Bnl/FGF signal transduction, and do not develop a normal ASP (2, 4, 18, 19).

The basic outlines of cytoneme-mediated and neuronal signaling are similar - both mechanisms involve cell extensions that make contacts with target cells where signals are exchanged. Several proteins (e.g., Capricious, Neuroglian, Shibire) found to be essential for cytoneme-mediated signaling in the ASP have been previously implicated in neuronal synapse formation (20–23). In addition, it was recently reported that the inward rectifying potassium channel Irk2 is required for Dpp release by wing disc cells (24). These features invite the question whether there are more extensive and deeper homologies (25).

Three key attributes of the ASP - the physical separation between ASP cells that receive Dpp or Bnl/FGF and disc cells that produce them, the genetic tools that can separately and independently target the disc and ASP cells, and the robust experimental accessibility for isolation and imaging of the ASP and wing disc - makes the ASP system uniquely powerful for studies of cell-cell signaling. This investigation focuses on functional homologies between cytoneme synapses where signaling proteins transfer to receiving cells and neuronal synapses where neurotransmitters released by presynaptic cells are taken up by postsynaptic cells. We present genetic, histological and functional evidence showing that signature features in the presynaptic and postsynaptic compartments of neuronal synapses, including key components of these compartments, as well as Ca^2+^ influx are essential for cytoneme-mediated signaling in the ASP.

## Results

### Calcium transients in cells and cytonemes of the Drosophila ASP

The ASP of the Drosophila third instar larva has approximately 120 cells that form a structure with a narrow proximal stalk, a bulbous medial region and a rounded distal tip (Fig. 1A,B). The tubular ASP is not radially symmetric: the (“lower layer”) cells closest to the disc epithelium have a smaller circumference than (“upper layer”) cells that are not juxtaposed to the disc. Cells in the lower layer medial region activate Dpp signaling (and are positive for dad-GFP, a reporter of Dpp signal transduction; Fig. 1C); cells at the tip have active Bnl/FGF signaling (and are dpERK positive; Fig. 1C). Cytonemes that extend from the medial region contain the Dpp receptor and take up Dpp from the disc; cytonemes that extend from the tip contain the Bnl/FGF receptor and take up Bnl/FGF from the disc (Fig. 1A,B’) (4). Cytonemes that take up signaling proteins contact the disc at distances comparable to a synaptic cleft, as indicated by GRASP fluorescence (Fig. 1D, *) (2). The GRASP system, which presents two complementary extracellular fragments of GFP tethered to transmembrane domains and expressed on different cells, was developed to label synaptic contacts in C. elegans (26). GRASP fluorescence marks sites of stable, close juxtaposition (~20-40 nm). To investigate calcium signaling in the ASP, we expressed the genetically encoded calcium sensor GCaMP6 (27) and detected low GFP fluorescence in all ASP cells (Fig. 1E-E””; Movie S1). Most cells had low fluorescence, and ~1 cell/2 ASPs had brighter fluorescence that did not vary during an observation period of 8 minutes, but approximately six cells in every ASP (n=10) had transient flashes of brighter fluorescence that averaged approximately 35 seconds. The GCaMP6 fluorescent transients in different regions of the ASP were as follows: in the “proximal” stalk (10 cells), in the upper layer “medial” region between the stalk and tip (50 cells), upper layer tip (10 cells), the lower layer medial region (40 cells), and lower layer tip (9 cells) (Fig. 1F). The per cell frequencies of bright transients (ΔF/F ≥ 30%) in the lower layer cells of the tip and medial region were approximately the same (0.15, 0.18, respectively; n=10) (Fig. 1G); these cells are active for Bnl/FGF (tip) and Dpp signaling (medial region). In upper layer optical sections, cells with transient flashes at the tip (0.2) were approximately 3X more frequent on a per cell basis than in either the upper medial region (0.06) or stalk (0.06; n=10) (Figs. 1G, 2I). The upper medial and stalk cells are less active for Bnl/FGF and Dpp signaling than cells in either the tip or lower medial regions. These observations indicate that cells with calcium oscillations are more frequent in cells that are active for Dpp and Bnl/FGF signaling. Expression in the ASP of RNAi directed against calmodulin, a ubiquitous, calcium-binding protein, perturbed ASP morphogenesis (Fig. S1A), suggesting that calcium oscillations may have an essential role in the ASP cells that are active in signal transduction during ASP development.

**Figure 1.**
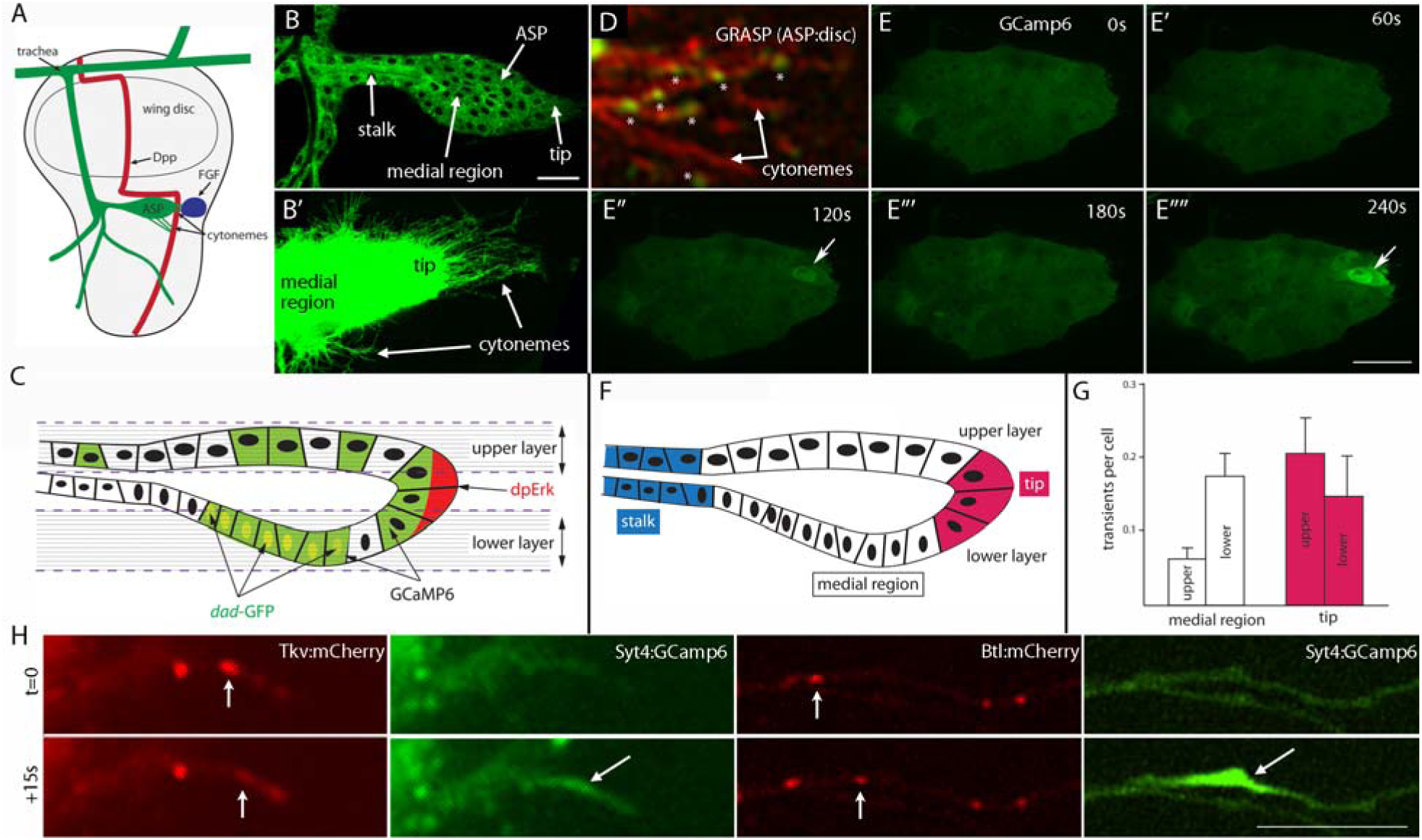
Calcium transients in the ASP and ASP cytonemes. (A) Cartoon showing a wing imaginal disc and associated trachea, with Dpp-(red) and FGF-expressing (blue) cells indicated. (B) Air sac primordium (ASP) expressing CD8:GFP from a late 3^rd^ instar larva. (B') Cytonemes labeled by CD8:GFP extending from ASP. (C) Cartoon showing sagittal view of ASP. Dashed lines represent approximate locations of the upper and lower optical sections. FGF-responsive cells are at the tip of ASP (dpErk), and Dpp-responsive cells are at the medial region *(dad-* GFP). (D) Contacts that ASP cytonemes (*, red) make with disc cells visualized by GRASP (green). (E-E””) Calcium transients in an ex vivo-cultured ASP expressing GCaMP6. Arrows point to the region of calcium increase. (F) Schematic drawing of an ASP marking stalk (blue), medial region (white) and tip (red). (G) Bar graph showing numbers of GCaMP6 transients/ASP cell. (H) Motile Tkv:mCherry and Bth:mCherry receptors (indicated by arrows) coinciding with calcium transient (green) in ASP cytonemes. Scale bars: 30μ μ **btl-Gal4 UAS-CD8:GFP/+;** *(D)* btl-Gal4 dpp-LHG/UAS-CD8:Cherry; UAS-CD4:GFP^1-10^ lexO-CD4:GFP^11^/+•, *(E-E””)* btl-Gal4 UAS-GCaMP6/+•, *(FI)* btl-Gal4/+; UAS-syt4:GCaMP6/UAS-Tkv:mCherry; *and* btl-Gal4 UAS-Btl:mCherry/+; UAS-syt4:GCaMP6/+.

Because cytonemes are conduits that traffic Dpp and Bnl/FGF from the disc to the ASP, we also characterized GCaMP6 fluorescence in ASP cytonemes. However, because the fluorescence of GCaMP6 in ASP cytonemes was undetectable (Fig. S2), we constructed a modified sensor that tethers GCaMP6 to the C-terminal cytoplasmic domain of the vesicular synaptic protein Synaptotagmin 4 (Syt4). The design was predicated on the idea that this domain of Syt4 might concentrate the chimeric GCaMP6 in cytonemes. We imaged preparations from animals that expressed Syt4:GCaMP6 as well as the Dpp receptor Tkv fused to mCherry (Tkv:mCherry; n=24) or the Bnl/FGF receptor Btl fused to mCherry (Btl:mCherry, n=33). Syt4:GCaMP6 expressed in the ASP generated uniform fluorescence that marked both cell bodies and cytonemes, and time-lapse imaging detected bright transients of GFP fluorescence in ASP cytonemes (average duration ~25 seconds; Fig. 1H; Movie S2, Movie S3). Every cytoneme marked by Syt4:GCaMP6 transients had an average of 2-3 transients during the period of observation (8 min, 5 sec exposure interval).

Motile puncta containing either Tkv and Btl are present in cytonemes that contact target disc cells, but have not been detected in cytonemes that are not in contact with disc cells (2). Similarly, calcium transients were observed only in cytonemes with motile Tkv- or FGFR-containing puncta, and not in cytonemes that lacked motile puncta. Although our images did not reveal how the transients relate to puncta motility or with ligand uptake, there was a clear correlation between calcium oscillations and movements of Dpp and Bnl/FGF receptors.

### Calcium oscillations in the ASP linked to cytoneme-mediated signaling

To investigate the basis for calcium transients in the ASP, we perturbed maintenance of intracellular Ca^2+^ by suppressing production of the endoplasmic reticulum ATPase SERCA, the main agent for Ca^2+^ uptake into the ER. We expressed an RNAi targeted to SERCA that had been used previously to study stress responses in wing discs (28). In this setting of reduced SERCA, ASP growth was inhibited, ASP morphogenesis was perturbed, and calcium transients in ASP cells were eliminated (0 transients ΔF/F≥0.3 in 8 ASPs; Movie S4). In addition, Dpp signal transduction in the ASP, which is essential for normal growth and morphogenesis and is cytoneme-dependent (2), was lowered (monitored by dad>GFP fluorescence; Fig. 2A-D, Fig. 5A), and the number of ASP cytonemes was suppressed. Because cytonemes extend and retract rapidly, and because the contacts they make are short-lived (6), this steady-state measure of cytoneme number does not distinguish between effects on rates of extension and retraction, or on contact duration.

**Figure 2.**
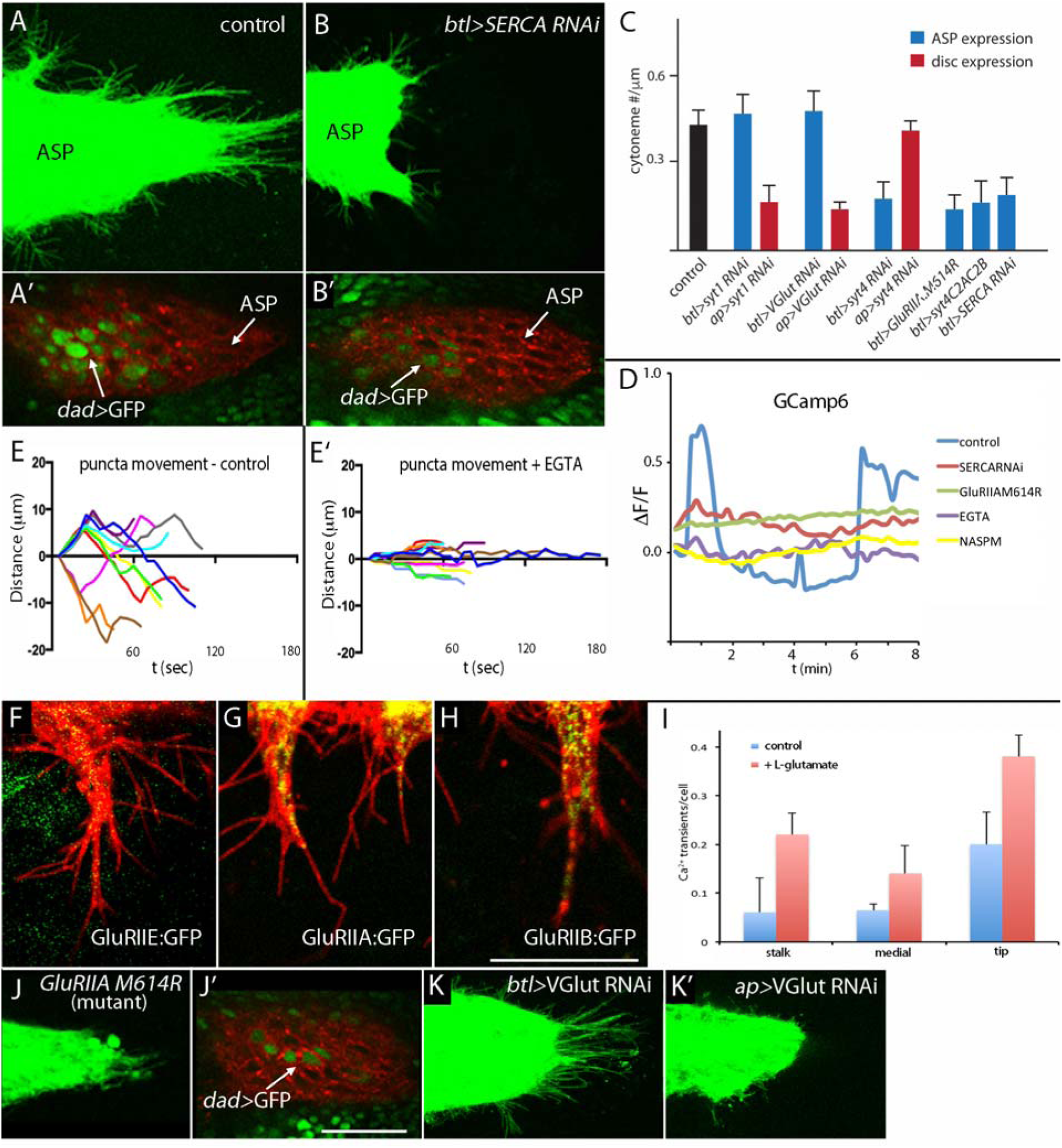
Cytoneme-mediated transport requires SERCA and Glutamate receptor (GluR). (A-B') Dependence of cytonemes-mediated Dpp signaling on SERCA. Downregulation of SERCA perturbed ASP development, and reduced the number of ASP cytonemes (A,B) and expression of *dad-GFP* (A',B'). (C) Bar graph plots the relative number of cytonemes in ASPs of control, *SERCARNAi, GluRIIA.M614R, sytlRNAi, syt4RNAi,* syt4 C2A C2B, and VGIutRNAi larvae; p values for difference between ASP and disc expression: p<0.001 (SERCARNAi), p<0.0001 (GluRIIA.M614R), p<0.001 (sytlRNAi), p<0.001 (syt4RNAi and syt4 C2A C2B), and p<0.0001 (VGIutRNAi). (D) Traces of Ca^2+^ transients in control, SERCA RNAi, GluRIIA.M614R, EGTA-treated, and NASPM-treated ASPs. (E,E') Kymographs depict the displacement often Tkv:mCherry puncta in control (E) and in presence of 2mM EGTA (E'). (F-H) Localization of GluRIIE:GFP (F), GluRIIA:GFP (G) and GluRIIB:GFP (H) in ASP cytonemes. (I) Graph representing the number of calcium transients/cell in the stalk, medial and tip regions of the upper layer of the ASP with standard deviation indicated; p values for difference between control and L-glutamate conditions: stalk (p<0.001), medial (p<0.01) and tip (p<0.001) regions. (J,J') Expression of *GluRIIA.M614R* reduced the number of ASP cytonemes and expression of cfad-GFP. (K,K') Expression of VGIutRNAi in the disc perturbed the ASP (K') but expression in the ASP did not (K). Scale bars: 15 μη (H); 30 μηη (J').

To investigate whether extracellular calcium is required for the Ca^2+^ transients, we reduced extracellular calcium with EGTA, a Ca^2+^ chelator. Incubation of ASP preparations in the presence of EGTA eliminated calcium transients in both ASP cells (0 transients ΔF/F≥0.3 in 5 ASPs) and ASP cytonemes (0 transients ΔF/F≥0.3 in cytonemes of 8 ASPs; Fig. 2D, Movie S4). In the absence of EGTA, Tkv:mCherry puncta moved in both anterograde and retrograde directions at approximately 0.41 μm/s, reaching distances of 10-20 μm (Fig. 2E, Movie S5). In the presence of EGTA (Fig. 2E’, Movie S6), Tkv:mCherry puncta had a more limited range of movement (all < 5 μm), suggesting that the motility of Tkv in cytonemes is dependent on uptake of extracellular Ca^2+^.

To investigate the process that generates Ca^2+^ transients, we expressed RNAi constructs directed against the transient receptor potential (Trp) and GluR calcium-permeable receptors. We targeted the thirteen identified Drosophila Trp family genes, one of which, inactive (iav), was shown previously to promote Ca^2+^ release from the endoplasmic reticulum and to be required for synapse development and neurotransmission (29). Expression in the ASP of RNAi directed against the thirteen Trp family genes did not affect ASP growth or ASP cytonemes (Fig. S1B-N). RNAi knockdown of the NMDA-type glutamate receptor was also without apparent effect (Fig. S1O). Although these results do not implicate these proteins in ASP signaling, the negative results do not rule out roles for Trp family or NMDA-type glutamate receptors. In contrast, we obtained genetic evidence that ASP cells require the non-NMDA ionotropic glutamate receptor (GluRII) (Fig. 2C).

GluRII is an AMPA/kainate-type receptor that has been implicated in retrograde signaling (30, 31). Isoforms of GluR have three common subunits (GluRIIC, GluRIID, GluRIIE) and a fourth subunit that is either GluRIIA or GluRIIB (32, 33). We examined GFP-tagged GluR subunits to monitor GluR presence in the ASP, characterizing a GluRIIE construct expressed by a fosmid transgene, and GluRIIA:GFP and GluRIIB:GFP that were ectopically over-expressed. Fluorescence of GluRIIE:GFP, GluRIIA:GFP and GluRIIB:GFP was detected in ASPs and in ASP cytonemes (Fig. 2F-H).

GluRIIA.M614R is a channel dead GluR subunit that acts as a dominant negative receptor that decreases quantal size of excitatory junctional potentials (33). The presence of GluRIIA.M614R in the ASP reduced the number of ASP cytonemes, eliminated calcium transients in the ASP (0 transients ΔF/F≥0.3 in 8 ASPs, Movie S4), perturbed ASP growth and morphogenesis, and reduced Dpp signaling and numbers of cytonemes (Fig. 2A,A’,C,J,J’, Fig. 5A). The presence of 1-Naphthylacetyl spermine trihydrochloride (NASPM), a pharmacological inhibitor of GluR (34), also eliminated ASP calcium transients (0 transients ΔF/F≥0.3 in 5 ASPs, Fig. 2D and Movie S4). In contrast, activation of GluR by the addition of L-glutamate (1mM) increased the number of ASP calcium transients (13 transients per ASP; n=5). The increase in calcium transient number affected all regions, including the stalk, upper layer medial region and tip, and was not restricted to the lower layer or tip that have active calcium transients in control conditions (Fig. 2I and Movie S7). Exogenously added L-glutamate increased both the amplitude approximately 3.2X and frequency of calcium transients approximately 2.2X (see also Fig. 5K,L)

The role of the glutamate receptor in the ASP suggests that the presynaptic wing disc cell might release glutamate. To test this prediction, we reduced expression of the vesicular glutamate transporter (VGlut) in either disc cells or in ASP cells (Fig. 2K,K’). Whereas reducing VGlut in the ASP had no apparent effect on ASP development or on the number of ASP cytonemes, targeting VGlut in the disc reduced the number of ASP cytonemes and resulted in abnormal ASP development (Fig. 2C,K,K’). Together with the observations that calcium transients in the ASP increased in response to L-glutamate and decreased in the presence of the GluR inhibitor NASPM, these data support the glutamatergic nature of the cytoneme synapse.

To control for possible non-specific effects on cell vitality and viability under conditions of compromised SERCA and GluRII function, the numbers of dividing and apoptotic cells were monitored in the presence of SERCA RNAi and GluRIIA.M614R. No apparent effects on frequency of mitosis or apoptosis were detected (Fig. S3). These observations suggest that cytoneme-mediated signaling involves intracellular calcium fluxes from both ER and extracellular sources.

### Postsynaptic functionality of Synaptotagmin 4 and Neuroligin 2 in cytoneme-mediated signaling

Vesicle fusion and receptor internalization in presynaptic and postsynaptic compartments involves Synaptotagmin Ca^2+^ binding proteins. In Drosophila, the Syt 1 isoform is the sensor for presynaptic vesicle exocytosis (35, 36), and Syt4 in postsynaptic compartments is involved in retrograde signaling (37). To investigate the role of Syt4 for cytoneme-mediated signaling, we expressed a Syt4-specific RNAi construct to determine if it is required at cytoneme synapses, and if so, if it is specific for the postsynaptic compartment. Whereas expression of Syt4 RNAi in the wing disc did not affect ASP development (Fig. 3A), expression of Syt4 RNAi in the ASP reduced Dpp uptake and signal transduction, perturbed development in the ASP, and decreased the number of ASP cytonemes (Figs. 2C, 3B, 5A and Fig. S4). ASP expression of a mutant Syt4 that does not bind Ca2+ (38) reduced Dpp signal transduction and perturbed development in the ASP (Fig. 3C); and in the viable Syt4^BA1^ null mutant (39), adult dorsal air sacs were reduced in size and malformed (Fig. S5). These observations suggest that Syt4 functions specifically in the ASP and is necessary for Dpp signaling in the ASP.

**Figure 3.**
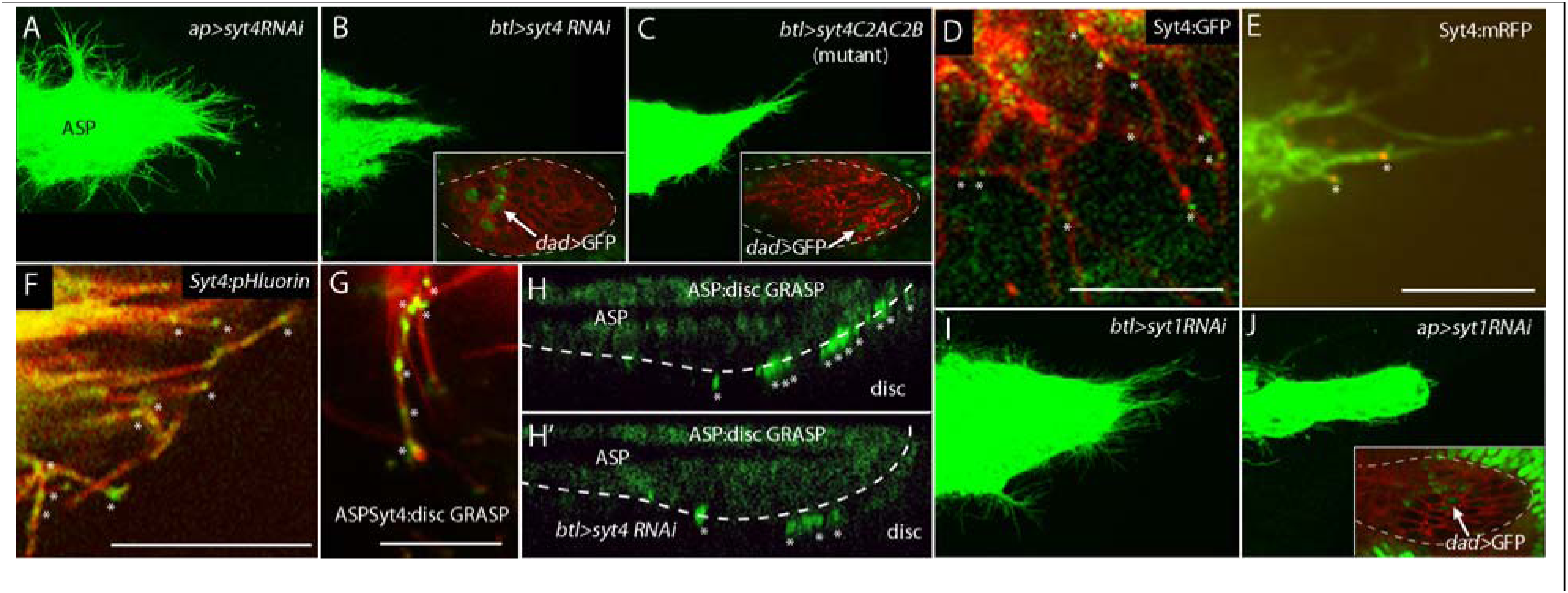
Cytoneme-mediated transport requires Synaptotagmin-4. (A,I) Syt1 in ASP or Syt4 in disc is dispensable for the ASP. (B,J) The ASP depends on Syt1 in wing disc and Syt4 in ASP. ASPs marked by membrane-tethered GFP and expressing RNAi constructs in either the ASP (driven by *btl-Gal4)* or dorsal compartment of the wing disc (driven by *ap-Gal4).* (B,C,J) ASP cytonemes and expression of *dad*-GFP (insets, arrows) were reduced with expression of syt1-RNAi in the disc, and syt4-RNAi or a mutant form of Syt4 that lacked Ca^2+^-binding sites in the ASP. *dad*-GFP analysis was in preparations expressing RNAi for 24 hours (by repressing Gal80^ts^ conditionally). (D) Syt4:GFP in ASP cytonemes. * indicates Syt4:GFP puncta. (E) An image frame from Movie S9 showing localization of Syt4:mRFP in ASP cytonemes. * indicates Syt4:mRFP puncta. (F) Syt4-pFllourin fusion protein expressed in the ASP. * indicates fluorescence of Syt4-pFllourin in ASP cytonemes. (G) GRASP fluorescence with Syt4:GFP^1-10^ in the ASP and GFP^11^ in φρ-expressing wing disc cells. * indicates syt4GRASP fluorescence in ASP cytonemes. (H,H') Sagittal images showing GRASP fluorescence at contacts (*) between ASP cytonemes and Dpp-expressing cells. ASPs outlined by dashed white lines; Scale bars: 15 μm (E); 10 μm (D), (F), (G).

To determine if Syt4 is in cytonemes, we examined several engineered forms of Syt4. Syt4:GFP is a fusion protein generated by CRISPR-mediated insertion into the endogenous Syt4 gene; it has wild type function and is expressed at endogenous levels (40). ASP cytonemes with Syt4:GFP and marked with CD8:mCherry had multiple fluorescent GFP puncta (Fig. 3D). Expression of a Syt4:mRFP fusion protein (btl-Gal4 UAS-Syt4:mRFP) also revealed the presence of Syt4 puncta in ASP cytonemes (Fig. 3E). The Syt4:mRFP puncta were motile (Movie S9).

To investigate the topology of Syt4 in the cytoneme membrane, we expressed Syt4-pHluorin, a Syt4 chimera tagged with a pH-sensitive variant GFP (38). pHluorin fluorescence is not detectable at the low pH of the interior of intracellular vesicles, but increases approximately 20x at neutral pH, and is detectable when exposed to the extracellular environment. Expression of Syt4-pHluorin in the ASP produced spots of bright GFP fluorescence along ASP cytonemes (Fig. 3F), suggesting that Syt4 in cytonemes is likely to be membrane-associated and exposed to the extracellular space.

The spots with Syt4-pHluorin fluorescence were immobile, in contrast to the cytoneme-associated puncta containing fluorescent Tkv or Btl (Movies S2, S3). This suggests that Syt4-rich spots might be sites where Syt4-containing vesicles are delivered. To characterize these sites further, we constructed a Syt4:GFP chimera and applied the GRASP (GFP Reconstitution Across Synaptic Partners) technique (26). GRASP can identify contacts that ASP cytonemes make with cells that express GFP, the peptide that complements GFP (25). Expression of Syt4:GFP in the ASP and of CD4:GFP in the dpp expression domain of the wing disc produced GFP fluorescence at discrete spots along ASP cytonemes (Fig. 3G). This pattern of GRASP fluorescence is similar to the pattern of Syt4:pHlourin fluorescence and is consistent with the idea that the Syt4 extracellular domain is exposed. It is also consistent with the idea that ASP cytonemes make multiple contacts with disc cells. A previous report described multiple contacts between cytonemes that extend from cells of the wing disc that produce the Hh signaling protein and wing disc cells that receive Hh (7).

To determine whether Syt4 is required for the contacts that ASP cytonemes make with disc cells, we compared the GRASP GFP fluorescence associated with ASP cytonemes in animals with normal or reduced levels of Syt4. In controls, expression of CD8:GFP in the ASP and of CD8:GFP in the dpp expression domain of the disc generated bright GFP fluorescence concentrated in the area between the lower layer of the ASP and the disc (Fig. 3H). Expression of syt4RNAi in the ASP significantly reduced GRASP fluorescence (Fig. 3H’). This result is similar to the effect previously observed for mutants that lack normal Caps function in ASP cytonemes (2), and it suggests that Syt4 is also required to make cytoneme synapses.

Neuroligins are synaptic adhesions proteins that participate in neuronal synapse formation in vertebrates and Drosophila (41–44). Drosophila has four Neuroligin orthologs (Nlg1-4) (41). We tested Nlg1 and Nlg2, and observed that expression of Nlg2-RNAi in the ASP inhibited ASP development. In contrast, expression of Nlg2-RNAi in the wing disc was without apparent effect, and no ASP abnormalities were observed in response to ASP expression of RNAi directed against Nlg1 (Fig. S1R,S,T). These results suggest that Nlg2 is required in the ASP for cytoneme-mediated signaling.

### Synaptic functions in signal protein-producing wing disc cells

To investigate whether functions that are required in the presynaptic compartment of neuronal synapses also play roles in cytoneme-dependent signaling, we analyzed animals defective for Syt 1, Synaptobrevin (Syb/dVAMP) and for the Cacophony (Cac; α1) and Straightjacket (Stj; α2δ) subunits of the primary presynaptic voltage-gated calcium channel. To investigate the role of Syt 1 in cytoneme-mediated signaling, we expressed a Syt 1-specific RNAi construct (45). Whereas Syt 1 RNAi expression in the ASP had no apparent effect, its expression in the wing disc reduced Dpp signal transduction in the ASP and resulted in abnormal ASP development (Figs. 2C, 3I,J, 5A). qPCR analysis (Fig. S6) indicated that syt 1 is expressed in the wing disc. These findings suggest that Syt 1 is required specifically in the presynaptic compartment of the cytoneme synapse.

Vertebrate Syb is a R-SNARE family component of neurotransmitter-containing vesicles that mediates the rapid docking response to inflow of Ca^2+^ ions in the active zone of presynaptic compartments (46). Drosophila encodes two homologs, one of which (n-Syb) is specific to neurons (47). n-Syb is functionally interchangeable with the other (Syb) (48), which is expressed ubiquitously and has a general role in membrane trafficking (Chin et al., 1993). In the wing pouch primordium of the wing disc, Syb is required for Wingless signaling (49) and by the cytonemes of the Hh-producing cells of the wing disc (8). Wing disc expression of the sybRNAi used by Yamazaki et al (49) stunted ASP growth and reduced Dpp signal transduction, and reduced the number of ASP cytonemes (Figs. 4A-C, 5A). In contrast, expression of this sybRNAi in the ASP was without apparent effect (Fig. 4D,D’). These results show that the defects caused by the sybRNAi line are tissue-specific, and suggest that the requirement for Syb function is specific to the presynaptic compartment of the cytoneme-wing disc synapse, a property that recalls the presynaptic role of n-Syb in neurons.

**Figure 4.**
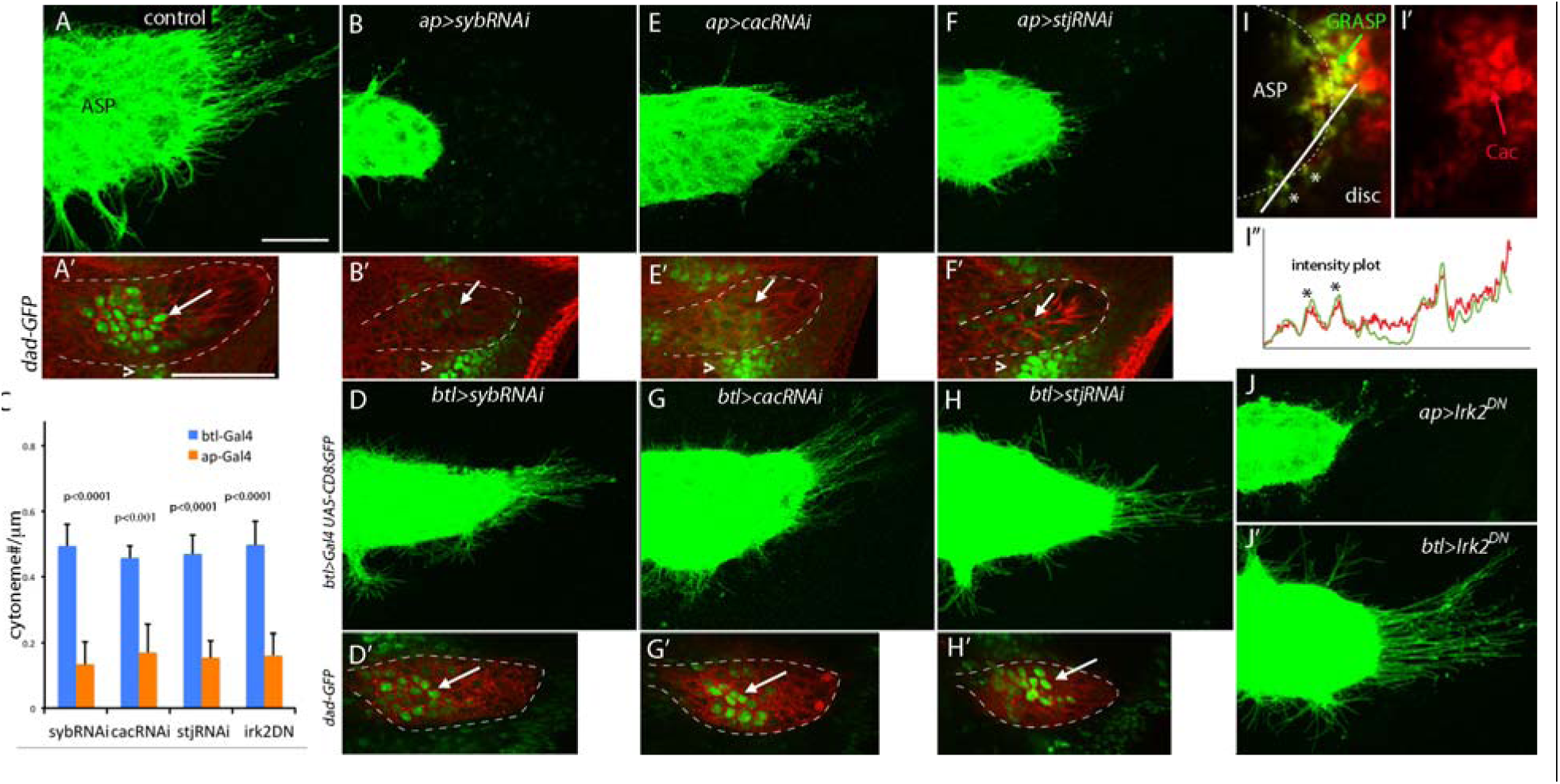
The ASP depends on the voltage-gated calcium channel and Synaptobrevin SNARE in the wing disc. (A-C,E-F') Downregulation in the wing disc of the Cac and Stj subunits of the voltage-gated calcium channel and of Syb reduced the number of ASP cytonemes and expression of*dad*-GFP. (C) Bar graph plots the relative number of cytonemes in ASPs under conditions of sybRNAi, cacRNAi, stjRNAi, and irk2^DN^ expression in the wing disc *(ap-Gal4)* or ASP (*btl-Gal4*). p values for difference between disc and ASP expression for each genotype. (D,D',G-H') Expression of cacRNAi, stjRNAi or sybRNAi in the ASP did not affect the ASP or cytoneme-mediated Dpp signal transduction. Arrows point to the dad-GFP expression in the ASP and arrowheads point to its expression in the μm (A), 50 μm (A').(I-I'') Cac:TdTomato and GRASP fluorescence coincide at contacts between ASP cytonemes and FGF-expressing wing disc cells. (I) Merged GFP + TdTomato fluorescence. ASP tip outlined with dashed white line, * indicates coinciding points of GRASP and TdTomato fluorescence; (I') TdTomato fluorescence; (I") intensity plot generated with ImageJ of GFP and TdTomato fluorescence in 3 pixel wide stripe (white line in I), normalized to equivalent minimum and maximum values. (J,J') Expression of *irk2^DN^* in the disc perturbed the ASP (J') but expression in the ASP did not (J)

**Figure 5.**
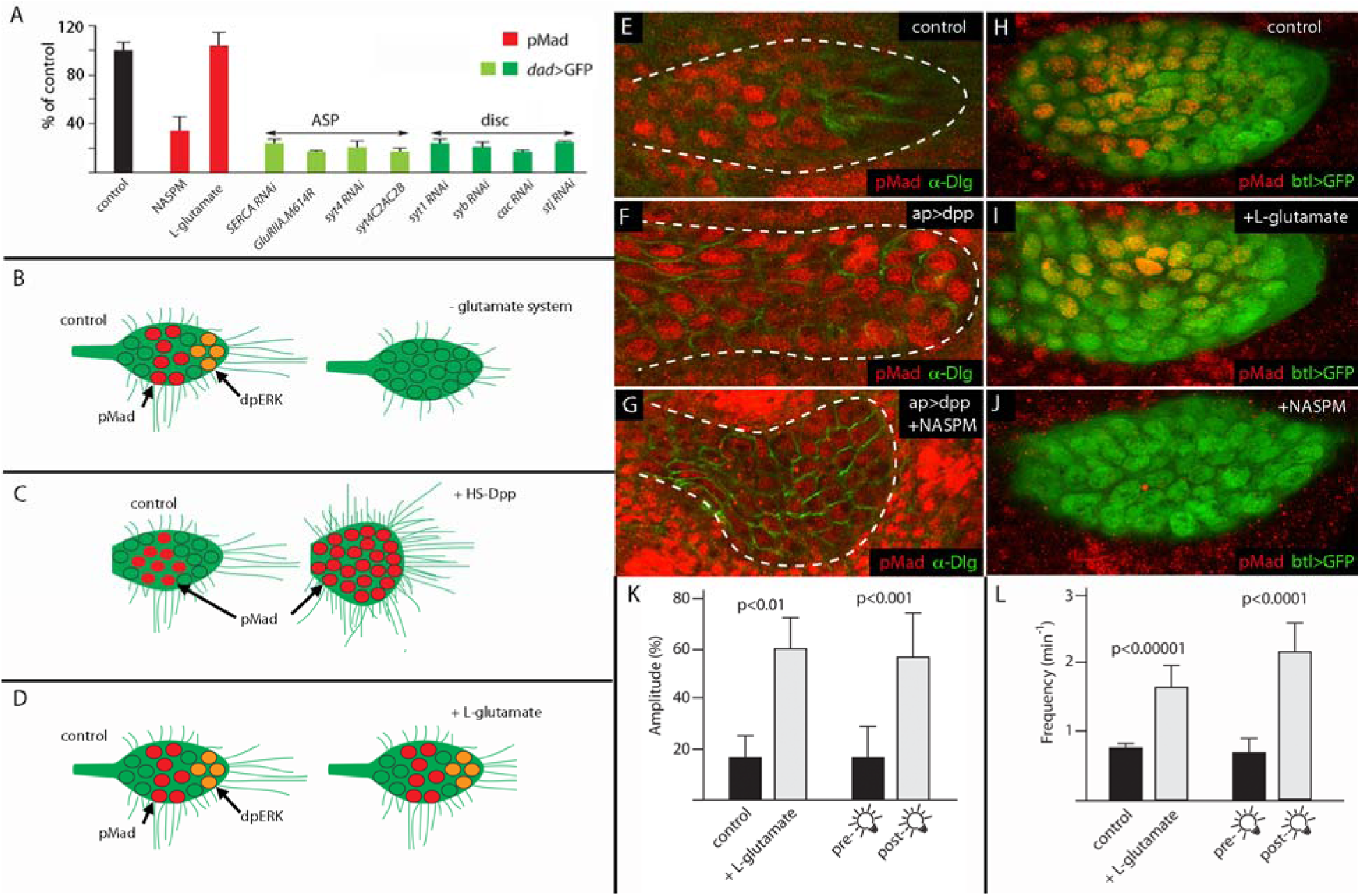
Dependence of Dpp signaling on glutamate and synaptic proteins. (A) Bar graph showing levels of Dpp signal transduction in ASP, fold changes relative to respective control. For chemical treatment: *btl-Gal4; UAS-GFP;* for ASP expression: *btl-Gal4 UAS-CD8:Cherry/tub-Gal80^ts^; dad-GFP*/+; for disc expression: *ap-Gal4/+; dad-GFP/+.* p values differences between control and experimental condition: NASPM (2.9E-03), L-glutamate (0.58*),SERCARNAi* (3.9E-06), *GluRIIA.M614R* (2.0E-07), *syt4RNAi* (2.4E-06), *syt4 C2AC2B* (2.9E-07), *sytlRNAi* (2.1E-06), *sybRNAi* (1.8E-06), *cacRNAi* (4.1E-07), *stjRNAi* (9.8E-07). (B,C,D) Drawings depicting changes of ASP cytonemes, signaling and morphology in response to (B) glutamate system inhibition, (C) ectopic Dpp over-expression (4), (D) addition of L-glutamate. (E-G) Increased Dpp signal transduction (pMad) in the ASP in response to ectopic Dpp expression in the disc (F) was blocked by presence of NASPM (G); (see Methods for protocol for conditional expression of *dpp).* (H-J) Dpp signal transduction in ASP preparations incubated for 3 hours in mock (H), in 1 mM L-glutamate (I), or in 10 μ frequency (L) of Ca^2+^ transients.

An essential component of the neuronal presynaptic compartment is the voltage-gated calcium channel (VGCC), which initiates neurotransmitter exocytosis in response to membrane depolarization. The Drosophila genes straightjacket (stj) and cacophony (cac) encode two of four subunits (49, 50). We determined that cac and stj are expressed in the wing disc by qPCR analysis (Fig. S6), and observed that expression in the wing disc of RNAi constructs targeting cac and stj transcripts changed ASP morphology, reduced Dpp signal transduction in the ASP, and reduced the numbers of ASP cytonemes (Fig. 4C, E-F’, 5A). These cac and stj RNAi constructs have been previously characterized and validated in several behavioral assays (51, 52). Expression of these RNAi constructs in the ASP was without apparent effect (Fig. 4G-H’), suggesting that Dpp signaling to the ASP requires VGCC specifically in the wing disc.

To determine if the VGCC is present at ASP-wing disc synaptic contacts, we expressed a functional, fluorescent-tagged Cac transgene in the dpp-expressing cells of wing disc together with GRASP constructs that marked the cytoneme synapses. Previous studies of this tagged Cac protein showed that it localized to the presynaptic active zone of developing neurons (53). We observed that expression of this tagged Cac protein in the wing disc revealed that Cac was present at sites of contact between ASP cytonemes and wing disc cells (Fig. 4I-I”). These results and the fact that suppression of stj transcripts by RNAi (Fig. 4F) caused phenotypes that were similar to the stj mutant (Fig. S1U), are consistent with the idea that the VGCC functions at the presynaptic compartment of wing disc cells for Dpp signaling from the wing disc to the ASP. Given the role of the VGCC at the neuronal synapse, these results suggest that the cytoneme synapse is sensitive to and activated by membrane depolarization.

We also investigated the inward rectifying potassium channel Irk2 for a role in signaling by the wing disc. Irk2 was shown previously to regulate Dpp release from wing disc cells and for wing disc development, and to participate in generation of calcium transients in the wing disc (24). Consistent with earlier studies (28), we detected calcium transients in the wing disc (Movie S10), and we observed that expression of a dominant negative mutant Irk2 in the wing disc affected ASP development (Fig. 4J) in ways that are consistent with inadequate Dpp signaling. In contrast, expression of Irk2 in the ASP had no apparent effect (Fig. 4J’).

### An essential role for glutamate in cytoneme-mediated signaling

Figure 5A summarizes our results showing that levels of Dpp signal transduction in the ASP are sensitive to suppression of the VGCC and Syb in the Dpp-producing disc cells and to suppression of SERCA, GluRII, and Syt4 in the ASP. In a late 3^rd^ instar larval stage ASP, cells with highest levels of Bnl/FGF signaling are at the tip and cells with highest levels of Dpp signaling are in the medial region (Fig. 1C, 5B,E,H). Ubiquitous ectopic expression of Dpp induces Dpp signaling in cells throughout the ASP, and cytonemes extend outwards without apparent directional bias and in greater numbers, and the ASP has a more bulbous form (Fig. 5C,F) (4). Because our genetic tests showed that the components of glutamatergic neuronal synapses are necessary for Dpp signaling in the ASP, we tested directly for response to glutamate and to NASPM. Whereas expression of Dpp throughout the region of disc near the ASP induced uniform, high level activation of Dpp signaling in the ASP (Fig. 5F), the presence of L-glutamate, which induced repetitive Ca^2+^ transients throughout the ASP (Fig. 5K,L and Movie S7), did not change the level or pattern of Dpp signaling in the ASP (Fig. 5A,D,H,I), and the presence of NASPM blocked Dpp signaling (Fig. 5A,B,G,J). NASPM did not reduce the number of ASP cytonemes (Fig. S7). These results show that the contribution of synaptic glutamate signaling is essential but not sufficient for cytoneme-mediated Dpp signaling, and that the glutamate system is essential for cytoneme function. The presence of (ineffective) cytonemes under conditions of NASPM inhibition contrasts with the reductions in numbers of cytonemes observed in mutant genotypes that also impair cytoneme-mediated signaling. However, because effects on transient structures under conditions of acute treatment (i.e. pharmacological inhibition) may not be comparable to effects after development under genetic impairment, the different outcomes of these treatments on numbers of cytonemes may not be indicative of mechanistic differences.

### Transmission at the cytoneme synapse

Basic structural features of the contact between the cytoneme tip and target cell – the terminus of a cellular extension, the constitution with common components, the gap of 20-40 nm – are shared with the neuronal synapse. In addition, wing disc cells that are targets of ASP cytonemes express proteins including the VGCC that also contribute essential functions to the transmission of signals from the presynaptic compartment of the neuronal synapse. Moreover, these functions are necessary for signaling from the wing disc to the ASP. To test directly whether this cytoneme-mediated signaling is induced by depolarization, we monitored calcium transients in ASP preparations dissected from animals that expressed a channelrhodopsin that functions as a light-gated ion channel (54). Wing disc-specific expression of the channelrhodopsin and photostimulation induced calcium transients in the ASP (Fig. 5K,L and Movie S8). The increased amplitude and frequency of the optogenetically-induced transients were similar to the observed stimulation by L-glutamate. We conclude that the calcium transients in the ASP are responses to trans-synaptic transmission.

## Discussion

In earlier work, we found that signaling between cells of the wing disc and ASP is cytoneme-mediated and involves the intercellular transfer of signaling proteins at synaptic contacts (2). Cytoneme synapses were characterized by GRASP fluorescence (2, 7), a technique that marks sites of close (approximately 20-40 nm) cell-cell apposition and that was developed to identify neuronal synapses (26). Cytoneme synapses are dependent on the transsynaptic adhesion proteins Capricious and Neuroglian that also have essential roles in neuronal synapse formation (20, 21, 55, 56). The work reported here presents genetic, histological and functional evidence that adds to the number of components and features that are common to both cytoneme and neuronal synapses. This work is the first to show that voltage-activated glutamate transmission is an essential element of signaling between epithelial cells.

We examined more than thirty genetic and pharmacological conditions to target Dpp signaling in the ASP. Because most of the genes are essential and null mutants are lethal early in development, the genetic conditions we characterized were partial loss-of-function - either in a hypomorphic mutant or generated by tissue-specific expression of partially defective proteins or gene-specific RNAi constructs. Expression conditions were adjusted to have no discernable effects on cell viability or cell division (Fig. S3). Partial loss-of-function conditions in the wing disc that targeted syt1, syb, cac, stj, Irk2, VGlut, or *Atpα*, all essential components of presynaptic neuronal compartments, reduced signaling in the ASP. In contrast, targeting these genes in the ASP had no apparent phenotype. Partial loss-of-function conditions in the ASP that targeted GluRII, syt4, or SERCA, all essential components of postsynaptic neuronal compartments, reduced signaling in the ASP; but targeting these genes in the wing disc had no effect on the ASP. This specificity, which implicates presynaptic glutamatergic functions only in the signal producing cells of the wing disc and postsynaptic glutamatergic functions only in the signal receiving cells of the ASP, is strong genetic evidence that the observed phenotypes are correctly attributed to the gene targets.

Histological characterizations showed that GluRII is present in cytonemes that generate the postsynaptic compartments (Fig. 2F-H), that the VGCC is present presynaptically at the cytoneme synapse, and that Syt4 is exposed on the plasma membrane of postsynaptic cytonemes (Fig. 3E-G) (38). These findings show that these key components of the glutamatergic synapse are segregated in similar ways at both cytoneme and neuronal synapses. The fact that optogenetic stimulation of a channelrhodopsin expressed specifically in the wing disc induced Ca^2+^ transients in the ASP is strong evidence of synaptic neurotransmitter signaling. We conclude that these epithelial cells make functional glutamatergic synapses.

The presence of glutamatergic synapses in non-neuronal, epithelial cells of the Drosophila wing disc and ASP is unexpected and without precedent, but is consistent with a comparative analysis of the GluR gene family that concluded that GluR signaling is present even in animals that lack a nervous system, such as Trichoplax (57). Plants also regulate Ca^2+^ influx with GluR-like channels that are required for growth and cell-cell communication (58, 59).

Our study implicates the glutamate neurotransmitter system in signaling by proteins such as Dpp at the cytoneme synapse, as inhibition of the glutamate receptor and genetic deficits of components of glutamate signaling also blocked Dpp signaling. This role of glutamate signaling at the cytoneme synpase should be understood in the context of specificity. Whereas ectopic over-expression of Dpp activates Dpp signaling ectopically, the presence of exogenous L-glutamate, which induces ectopic and unusually frequent Ca^2+^ transients, does not increase Dpp signaling. Activation of the glutamate receptor system is therefore not sufficient to activate the signal transduction pathway. Cells that express multiple types of signal protein receptors are competent to respond to the signal proteins - in the case of the ASP cells, which express both the Dpp and Bnl/FGF receptors, to bind their specific ligands and activate the specific downstream signal transduction pathway. Although the glutamate receptor system is essential for the activation of both signal transduction pathways, signaling is specific to different signal proteins because it is dependent on binding of a signal protein with its receptor.

Studies of the glutamatergic components at neuronal synapses show that synapse structure is normal in their absence, but signaling across the synapse is defective (37, 38, 60, 61). Although we might suggest that cytoneme synapses provide a function that is analogous to the neuronal synapses – a setting in which signal release is regulated in the context of uptake - our understanding of the processes that produce cytonemes and cytoneme synapses, or release Dpp from the transmitting cell and take it up postsynaptically, is too rudimentary to entertain models for the precise role of glutamate signaling at a cytoneme synapse. In contrast to neuronal synapses, cytoneme synapses are transient and our studies of cytonemes and cytoneme-synapses have been limited to steady-state measures. Consequently, our studies do not distinguish whether glutamate signaling contributes to cytoneme production, or if by facilitating signal release and uptake, it contributes to stabilization. Studies in both vertebrates and invertebrates implicate morphogen signaling proteins (e.g., Wnt/Wg, TGF-11BMP/Dpp, Hh, EGF) in neural development and neuron function (62–69). Ex vivo and in vivo experiments identify roles for signaling proteins in neuronal polarity, axon pathfinding, synaptogenesis, synaptic plasticity and long-term potentiation (70). Linkage of these roles of signaling proteins to mechanisms of synaptic excitability has not been considered, nor has synaptic excitability been associated with morphogen signaling. Our discovery of the essential role of the glutamate receptor system in morphogen signaling suggests that the linkage may be direct.

## Supporting information

## Acknowledgements

We thank: Drs. T. Littleton, D. Volfson, M. O’Connor, E. Martin-Blanco, K. Scott, E. Serpe, D. Anderson, R. Ordway, and the Vienna Drosophila RNAi Center, Kyoto Stock Center, and Bloomington Stock Center for fly stocks; T. Littleton and D. Volfson for syt4-mRFP construct; V. Ruta for the sytGCaMP6s plasmid; S. Roy for his help and advice; all members of Kornberg lab for discussion and constructive suggestions. Funding: NIH grants R01GM030637 and R35GM122548 to T.B.K. and 5T32HL007731 to H.H.. Author contributions: conceptualization - HH, TBK; investigation - HH, SL; Writing - HH, TBK. Competing interests: none.

## References

1 M. Sato, T. B. Kornberg, FGF is an essential mitogen and chemoattractant for the air sacs of the Drosophila tracheal system. Dev. Cell. 3, 195–207 (2002).

2 S. Roy, H. Huang, S. Liu, T. B. Kornberg, Cytoneme-Mediated Contact-Dependent Transport of the Drosophila Decapentaplegic Signaling Protein. Science. 343, 1244624–1244624 (2014).

3 C. F. A. Ramirez-Weber, T. B. Kornberg, Cytonemes: cellular processes that project to the principal signaling center in Drosophila imaginal discs. Cell. 97, 599–607 (1999).

4 S. Roy, F. Hsiung, T. B. Kornberg, Specificity of Drosophila cytonemes for distinct signaling pathways. Science. 332, 354–358 (2011).

5 F. Hsiung, F. A. Ramirez-Weber, D. D. Iwaki, T. B. Kornberg, Dependence of Drosophila wing imaginal disc cytonemes on Decapentaplegic. Nature. 437, 560–563 (2005).

6 M. Bischoff et al., Cytonemes are required for the establishment of a normal Hedgehog morphogen gradient in Drosophila epithelia. Nat Cell Biol. 15, 1269–1281 (2013).

7 L. González-Méndez, I. Seijo-Barandiarán, I. Guerrero, Cytoneme-mediated cell-cell contacts for hedgehog reception. Elife. 6 (2017), doi:10.7554/eLife.24045.

8 W. Chen, H. Huang, R. Hatori, T. B. Kornberg, Essential basal cytonemes take up Hedgehog in the Drosophila wing imaginal disc. Development (2017), doi:10.1242/dev.149856.

9 M. Inaba, M. Buszczak, Y. M. Yamashita, Nanotubes mediate niche-stem-cell signalling in the Drosophila testis. Nature. 523, 329–332 (2015).

10 T. J. Fuwa, T. Kinoshita, H. Nishida, S. Nishihara, Reduction of T antigen causes loss of hematopoietic progenitors in Drosophila through the inhibition of filopodial extensions from the hematopoietic niche. Dev. Biol. 401, 206–219 (2015).

11 E. Stanganello et al., Filopodia-based Wnt transport during vertebrate tissue patterning. Nat. Commun. 6, 5846 (2015).

12 L. Caneparo, P. Pantazis, W. Dempsey, S. E. Fraser, Intercellular bridges in vertebrate gastrulation. PLoS One. 6 (2011), doi:10.1371/journal.pone.0020230.

13 D. S. Eom, E. J. Bain, L. B. Patterson, M. E. Grout, D. M. Parichy, Long-distance communication by specialized cellular projections during pigment pattern development and evolution. Elife. 4 (2015), doi:10.7554/eLife.12401.

14 T. A. Sanders, E. Llagostera, M. Barna, Specialized filopodia direct long-range transport of SHH during vertebrate tissue patterning. Nature. 497, 628–632 (2013).

15 J. C. Fierro-González, M. D. White, J. C. Silva, N. Plachta, Cadherin-dependent filopodia control preimplantation embryo compaction. Nat. Cell Biol. 15, 1424–1433 (2013).

16 J. C. Snyder et al., Lgr4 and Lgr5 drive the formation of long actin-rich cytoneme-like membrane protrusions. J. Cell Sci. 128, 1230–1240 (2015).

17 A. Callejo et al., Dispatched mediates Hedgehog basolateral release to form the long-range morphogenetic gradient in the Drosophila wing disk epithelium. Proc Natl Acad Sci U S A (2011), doi:10.1073/pnas.1106881108.

18 R. H. Huang, T. B. Kornberg, Myoblast cytonemes mediate Wg signaling from the wing imaginal disc and Delta-Notch signaling to the air sac primordium. Elife. 4, e06114 (2015).

19 H. Huang, T. B. Kornberg, Cells must express components of the planar cell polarity system and extracellular matrix to support cytonemes. Elife. 5 (2016), doi:10.7554/eLife.18979.

20 E. Shishido, M. Takeichi, A. Nose, Drosophila synapse formation: Regulation by transmembrane protein with leu-rich repeats, capricious. Science (80-.). 280 (1998), pp. 2118–2121.

21 H. Kohsaka, A. Nose, Target recognition at the tips of postsynaptic filopodia: accumulation and function of Capricious. Development. 136, 1127–1135 (2009).

22 H. Taniguchi, E. Shishido, M. Takeichi, A. Nose, Functional dissection of Drosophila Capricious: Its novel roles in neuronal pathfinding and selective synapse formation. J. Neurobiol. 42, 104–116 (2000).

23 W. Hong et al., Leucine-rich repeat transmembrane proteins instruct discrete dendrite targeting in an olfactory map. Nat. Neurosci. 12, 1542–1550 (2009).

24 G. R. Dahal, S. J. Pradhan, E. A. Bates, Inwardly rectifying potassium channels influence Drosophila wing morphogenesis by regulating Dpp release. Development. 144, 2771–2783 (2017).

25 T. B. Kornberg, S. Roy, Communicating by touch--neurons are not alone. Trends Cell Biol. 24, 370–376 (2014).

26 E. H. Feinberg et al., GFP Reconstitution Across Synaptic Partners (GRASP) defines cell contacts and synapses in living nervous systems. Neuron. 57, 353–363 (2008).

27 T.-W. Chen et al., Ultrasensitive fluorescent proteins for imaging neuronal activity. Nature. 499, 295–300 (2013).

28 S. Restrepo, K. Basler, Drosophila wing imaginal discs respond to mechanical injury via slow InsP3R-mediated intercellular calcium waves. Nat. Commun. 7, 12450 (2016).

29 C. O. Wong et al., A TRPV channel in drosophila motor neurons regulates presynaptic resting Ca2+ levels, synapse growth, and synaptic transmission. Neuron. 84, 764–777 (2014).

30 S. A. Petersen, R. D. Fetter, J. N. Noordermeer, C. S. Goodman, A. DiAntonio, Genetic analysis of glutamate receptors in drosophila reveals a retrograde signal regulating presynaptic transmitter release. Neuron. 19, 1237–1248 (1997).

31 C. M. Schuster et al., Molecular cloning of an invertebrate glutamate receptor subunit expressed in Drosophila muscle. Science (80-.). 254, 112–114 (1991).

32 G. Qin, Four Different Subunits Are Essential for Expressing the Synaptic Glutamate Receptor at Neuromuscular Junctions of Drosophila. J. Neurosci. 25, 3209–3218 (2005).

33 A. DiAntonio, S. A. Petersen, M. Heckmann, C. S. Goodman, Glutamate receptor expression regulates quantal size and quantal content at the Drosophila neuromuscular junction. J. Neurosci. 19, 3023–32 (1999).

34 M. Koike, M. Iino, S. Ozawa, Blocking effect of 1-naphthyl acetyl spermine on Ca(2+)-permeable AMPA receptors in cultured rat hippocampal neurons. Neurosci. Res. 29, 27–36 (1997).

35 J. T. Littleton, M. Stern, K. Schulze, M. Perin, H. J. Bellen, Mutational analysis of Drosophila synaptotagmin demonstrates its essential role in Ca2+-activated neurotransmitter release. Cell. 74, 1125–1134 (1993).

36 M. Geppert et al., Synaptotagmin I: A major Ca2+ sensor for transmitter release at a central synapse. Cell. 79, 717–727 (1994).

37 C. F. Barber, R. A. Jorquera, J. E. Melom, J. T. Littleton, Postsynaptic regulation of synaptic plasticity by synaptotagmin 4 requires both C2 domains. J. Cell Biol. 187, 295–310 (2009).

38 M. Yoshihara, B. Adolfsen, K. T. Galle, J. T. Littleton, Retrograde signaling by Syt 4 induces presynaptic release and synapse-specific growth. Science (80-.). 310, 858–863 (2005).

39 B. Adolfsen, S. Saraswati, M. Yoshihara, J. T. Littleton, Synaptotagmins are trafficked to distinct subcellular domains including the postsynaptic compartment. J. Cell Biol. 166, 249–260 (2004).

40 K. P. Harris, Y. V. Zhang, Z. D. Piccioli, N. Perrimon, J. Troy Littleton, The postsynaptic t-SNARE syntaxin 4 controls traffic of neuroligin 1 and synaptotagmin 4 to regulate retrograde signaling. Elife. 5 (2016), doi:10.7554/eLife.13881.

41 M. Sun et al., Neuroligin 2 is required for synapse development and function at the Drosophila neuromuscular junction. J. Neurosci. 31, 687–699 (2011).

42 Y.-C. Chen et al., Drosophila Neuroligin 2 is Required Presynaptically and Postsynaptically for Proper Synaptic Differentiation and Synaptic Transmission. J. Neurosci. 32, 16018–16030 (2012).

43 T. C. Südhof, Neuroligins and neurexins link synaptic function to cognitive disease. Nature. 455 (2008), pp. 903–911.

44 A. M. Craig, Y. Kang, Neurexin-neuroligin signaling in synapse development. Curr. Opin. Neurobiol. 17 (2007), pp. 43–52.

45 M. M. Paul et al., Bruchpilot and Synaptotagmin collaborate to drive rapid glutamate release and active zone differentiation. Front. Cell. Neurosci. 9 (2015), doi:10.3389/fncel.2015.00029.

46 Sudhof, Neurotransmitter release: the last milisecond in the life of a synaptic vesicle. Neuron. 80, 675–690 (2013).

47 A. DiAntonio et al., Identification and characterization of Drosophila genes for synaptic vesicle proteins. J. Neurosci. 13, 4924–4935 (1993).

48 S. Bhattacharya et al., Members of the synaptobrevin/vesicle-associated membrane protein (VAMP) family in Drosophila are functionally interchangeable in vivo for neurotransmitter release and cell viability. Proc. Natl. Acad. Sci. U. S. A. 99, 13867–13872 (2002).

49 Y. Yamazaki et al., Godzilla-dependent transcytosis promotes Wingless signalling in Drosophila wing imaginal discs. Nat. Cell Biol. 18, 451–457 (2016).

50 F. Kawasaki, S. C. Collins, R. W. Ordway, Synaptic calcium-channel function in Drosophila: analysis and transformation rescue of temperature-sensitive paralytic and lethal mutations of cacophony. J. Neurosci. 22, 5856–5864 (2002).

51 G. G. Neely et al., TrpA1 regulates thermal nociception in Drosophila. PLoS One. 6 (2011), doi:10.1371/journal.pone.0024343.

52 A. Saras, M. A. Tanouye, Mutations of the Calcium Channel Gene cacophony Suppress Seizures in Drosophila. PLoS Genet. 12 (2016), doi:10.1371/journal.pgen.1005784.

53 F. Kawasaki, Active Zone Localization of Presynaptic Calcium Channels Encoded by the cacophony Locus of Drosophila. J. Neurosci. 24, 282–285 (2004).

54 K. Watanabe et al., A Circuit Node that Integrates Convergent Input from Neuromodulatory and Social Behavior-Promoting Neurons to Control Aggression in Drosophila. Neuron. 95, 1112–1128.e7 (2017).

55 E. M. Enneking et al., Transsynaptic Coordination of Synaptic Growth, Function, and Stability by the L1-Type CAM Neuroglian. PLoS Biol. 11 (2013), doi:10.1371/journal.pbio.1001537.

56 M. Yamamoto, R. Ueda, K. Takahashi, K. Saigo, T. Uemura, Control of Axonal Sprouting and Dendrite Branching by the Nrg-Ank Complex at the Neuron-Glia Interface. Curr. Biol. 16, 1678–1683 (2006).

57 L. L. Moroz et al., The ctenophore genome and the evolutionary origins of neural systems. Nature. 510, 109–114 (2014).

58 M. M. Wudick et al., CORNICHON sorting and regulation of GLR channels underlie pollen tube Ca homeostasis. Science (80-.). 360, 533–536 (2018).

59 C. Ortiz-Ramírez et al., GLUTAMATE RECEPTOR-LIKE channels are essential for chemotaxis and reproduction in mosses. Nature. 549, 91–95 (2017).

60 C. V. Ly, C. K. Yao, P. Verstreken, T. Ohyama, H. J. Bellen, straightjacket is required for the synaptic stabilization of cacophony, a voltage-gated calcium channel α1 subunit. J. Cell Biol. 181, 157–170 (2008).

61 D. K. Dickman, P. T. Kurshan, T. L. Schwarz, Mutations in a Drosophila 2 Voltage-Gated Calcium Channel Subunit Reveal a Crucial Synaptic Function. J. Neurosci. (2008), doi:10.1523/JNEUROSCI.4498-07.2008.

62 Z. Huang, S. Kunes, Hedgehog, transmitted along retinal axons, triggers neurogenesis in the developing visual centers of the Drosophila brain. Cell. 86, 411–422 (1996).

63 C. Korkut et al., Trans-Synaptic Transmission of Vesicular Wnt Signals through Evi/Wntless. Cell. 139, 393–404 (2009).

64 S. Yogev, E. D. Schejter, B. Z. Shilo, Polarized secretion of drosophila EGFR ligand from photoreceptor neurons is controlled by ER localization of the ligand-processing machinery. PLoS Biol. 8 (2010), doi:10.1371/journal.pbio.1000505.

65 A. C. Hall, F. R. Lucas, P. C. Salinas, Axonal remodeling and synaptic differentiation in the cerebellum is regulated by WNT-7a signaling. Cell. 100, 525–535 (2000).

66 S. Yoshikawa, R. D. McKinnon, M. Kokel, J. B. Thomas, Wnt-mediated axon guidance via the Drosophila derailed receptor. Nature. 422, 583–588 (2003).

67 P. Ruth et al., The Drosophila BMP type II receptor Wishful Thinking regulates neuromuscular synapse morphology and function. J Neurophysiol. 16, 649–658 (2002).

68 M. Serpe, M. B. O’Connor, The metalloprotease Tolloid-related and its TGF--like substrate Dawdle regulate Drosophila motoneuron axon guidance. Development. 133, 4969–4979 (2006).

69 T. M. Jessell, Neuronal specification in the spinal cord: Inductive signals and transcriptional codes. Nat. Rev. Genet. 1 (2000), pp. 20–29.

70 M. Park, K. Shen, WNTs in synapse formation and neuronal circuitry. EMBO J. 31 (2012), pp. 2697–2704.

71 N. Ninov et al., Dpp Signaling Directs Cell Motility and Invasiveness during Epithelial Morphogenesis. Curr. Biol. 20, 513–520 (2010).

72 R. Cohn, I. Morantte, V. Ruta, Coordinated and Compartmentalized Neuromodulation Shapes Sensory Processing in Drosophila. Cell. 163, 1742–1755 (2015).

73 J. Zartman, S. Restrepo, K. Basler, A high-throughput template for optimizing Drosophila organ culture with response-surface methods. Development. 140, 667–674 (2013).

74 A. Guha, L. Lin, T. B. Kornberg, Regulation of Drosophila matrix metalloprotease Mmp2 is essential for wing imaginal disc:trachea association and air sac tubulogenesis. Dev. Biol. 335, 317–326 (2009).

