## Supplementary material for "Glutamate signaling at cytoneme synapses"

**Supplementary Materials:**

*Drosophila* stocks and husbandry

Flies were reared on standard cornmeal and agar medium at 22–25°C, unless otherwise stated. *btl*-Gal4, *UAS*-Tkv:mCherry and *UAS*-Btl:mCherry were described (*2*). Syt4:GFP, *UAS*-Syt4-S284A, *UAS*-Syt4-D292,427,429N, *UAS*-Syt4:mRFP, and *UAS*-Syt4-pHluorin, from T. Littleton (*37*, *38*, *40*); *UAS*-GluRIIA:GFP and *UAS*-GluRIIB:GFP from M. O’Connor; *dad*-GFP, from E. Martin-Blanco (*71*); *lexO*-CD4-GFP^11^ and *UAS*-CD4-GFP^1–10^ from K. Scott; RNAi targeting TRP channels and glutamate receptors and Fosmid lines from the Vienna Drosophila RNAi Center; *UAS*-Syt4GCaMP6, *UAS*-GCaMP6, *UAS*-SERCA RNAi, *UAS*-GluRIIA.M614R from the Bloomington Stock Center.

Transgenic flies

The design of syt4GCaMP6, based on syt1GCaMP6 (*72*) joined the coding sequence of GCaMP6 (Cohn et al., 2015) (Addgene) to the C-terminus of Syt4 (Barber et al., 2009) with an intervening 3XGS linker. The fragment (Syt4GCaMP6) was ligated to attB-containing pUAST. To generate the *UAS*-Syt4-GFP^1-10^ transgene, the coding region of Syt4 was amplified from a *syt4* cDNA clone (from T. Littleton) and inserted into an attB site containing pUAST vector. GFP^1-10^ (Addgene) fragment was joined to the 5’ end of Syt4 and constructs were injected into PBac{y[+]-attP-3B}VK00033 recipient flies by Rainbow Transgenic Flies.

Histology

Wing imaginal discs and trachea were dissected in cold PBS and were mounted in a drop of PBS underneath a coverslip using the ‘hanging drop’ method (*2*). Samples were imaged with a Leica TCS SPE confocal or an Olympus FV3000 inverted confocal laser scanning microscope. For wing disc culture, wandering third instar larvae were dissected in PBS. Wing imaginal discs together with Tr2 tracheal branches were incubated in WM1 medium (*73*). To image the ASP, the columnar layer of wing disc was proximal to the coverslip. Images and videos were taken with a Nikon spinning-disc confocal microscope and processed with NIS-Elements. Adult dorsal air sacs were viewed in thoraces of newly enclosed, decapitated flies mounted in halocarbon oil, and imaged with a Leica Stereo Microscope, as described.(*74*).

Overexpression and ectopic expression

*tub-Gal80^ts^* was utilized to restrict transgene expression to early L3 stage. For expression, animals were cultured at 18°C until L3 and incubated at 29°C for 1 day before dissection. Under this regimen, cell proliferation, apoptosis and cell polarity were unaltered (see Fig. S3). *ap-Gal4* and *btl-Gal4* drivers were used to express *UAS-dpp*, *UAS-syt1RNAi*, *UAS-sybRNAi*, *UAS-cacRNAi*, *UAS-syt4RNAi*, *UAS-irk2^DN^*, *UAS-SERCARNAi*, *UAS-GluRIIA.M614R*.

Calcium imaging and Drosophila wing disc culture

Larvae were staged and dissected in PBS. Wing imaginal discs and associated trachea were mounted in chamber slides in hemolymph-like (HL) media (108 mM NaCl, 5 mM KCl, 2 mM CaCl2, 8.2 mM MgCl2, 4 mM NaHCO3, 1 mM NaH2PO4, 5 mM trehalose, 10 mM sucrose and 5 mM HEPES, pH 7.5) for imaging in the presence of EGTA, NASPM, or L-glutamate, or in Schneider’s Drosophila medium containing 5% fly extract, insulin, and penicillin-streptomycin for all other imaging studies. Because imaging different optical planes was necessary to monitor the upper and lower levels of the ASP, fluorescence was monitored separately in the other upper and lower layers. Movies were captured with a Nikon spinning-disc confocal microscope with 405nm, 488 nm, 561 nm or 640 nm wavelength lasers.

Optogenetic activation

Prior to dissection, larvae expressing the Chrimson channelrhodopsin (*54*) in the *ap* domain of the wing disc were grown for three days on standard medium supplemented with 400 l of 400 μM all-trans retinal. GCaMP6 (*btl*-LHG lexO-GCaMP6) fluorescence was monitored with a spinning-disc confocal microscope. Photostimulation was with a 640 nm laser for 10 sec.

Immunohistochemistry

Wing imaginal discs and trachea were fixed in 4% formaldehyde. After several washes, the samples were permeablized with TritonX-100, blocked in 10% donkey serum and followed by incubation with primary antibodies (α-pMad (Abcam), α-Dlg (Developmental Studies Hybridoma Bank)) and secondary antibodies conjugated to Alexa Fluor 405, 488, 555, or 647 (Jackson ImmunoResearch, West Grove, PA). Samples were mounted in Vectashield. Images were captured by an Olympus FV3000 inverted confocal laser scanning microscope.

Statistical analysis of calcium pulse

A region of interest (ROI) that encompassed the most active cell in the ASP was selected. The intensity of Green fluorescence was measured with ICY (http://icy.bioimageanalysis.org/) software and imageJ. Changes in GCaMP intensity of ROIs were calculated in Excel. The ΔF/F graph was plotted in GraphPad Prism.

Quantification of the kinetics of Tkv:Cherry

Motile Tkv:mCherry puncta were tracked using MtrackJ tool in ImageJ. The displacements of individual puncta were measured and used for the analysis. The kymograph was generated using GraphPad Prism.

qRT-PCR

The Zymo Research RNA MicroPrep kit (Cat. #R1060) was used to extract total RNA from 30 control wild type wing discs and 30 wing discs expressing *tub-Gal4 UAS-syt1RNAi*. Reverse transcription was carried out using the Applied Biosystem High Capacity RNA-to-cDNA (Cat. #4387406). qPCR reactions were performed with a BioRad C1000 Touch Thermal Cycler and SYBR Green (Bioline).

**Figure 1.** Genotypes: (B,B’) *btl-Gal4 UAS-CD8:GFP/+*; (D) *btl-Gal4 dpp-LHG/UAS-CD8:Cherry;* *UAS-CD4:GFP^1-10^ lexO-CD4:GFP^11^/+*; (E-E’’’’) *btl-Gal4 UAS-GCaMP6/+*; (H) *btl-Gal4/+ ; UAS-syt4:GCaMP6/UAS-Tkv:mCherry;* and *btl-Gal4 UAS-Btl:mCherry/+ ; UAS-syt4:GCaMP6/+*.

**Figure 2.** Genotypes: (A) *btl-Gal4 UAS-CD8GFP/+*; (A’) *btl-Gal4 UAS-CD8Cherry/Gal80^ts^; dad-GFP/+*; (B) *btl-Gal4 UAS-CD8GFP/+; UAS-SERCA RNAi/+*; (B’) *btl-Gal4 UAS-CD8Cherry/Gal80^ts^; dad-GFP/SERCARNAi*; (J) *btl-Gal4 UAS-CD8GFP/UAS-GluRIIA.M614R*; (J’) *btl-Gal4 UAS-CD8Cherry/UAS-GluRIIA.M614R; dad-GFP/Gal80^ts^*; (K) *btl-Gal4 UAS-CD8GFP*/+ ; *UAS-VGlut RNAi*/+; (K’) *ap-Gal4/+ ; btl-LHG lexO-CD2GFP/UAS-VGlut RNAi.*

**Figure 3.** Genotypes: (A) *ap-Gal4/+; btl-LHG lexO-CD2:GFP/UAS-syt4RNAi*; (B) *btl-Gal4 UAS-CD8GFP/+; UAS-syt4RNAi/+*; (inset) *btl-Gal4 UAS-CD8Cherry/Gal80^ts^; dad-GFP/syt4RNAi*; (C) *btl-Gal4 UAS-CD8GFP/+; UAS–syt4[C2A D4N, C2B D3,4N]/+*; (inset) *btl-Gal4 UAS-CD8Cherry/+; dad-GFP/UAS–syt4[C2A D4N, C2B D3,4N]*; (D) *btl-Gal4 UAS-CD8Cherry/+; syt4:GFP/+*; (E) *btl-Gal4 UAS-CD8GFP/+; UAS-syt4:mRFP/+*; (F) *btl-Gal4 UAS-CD8Cherry/+; UAS-Syt4-pHluorin/+*; (G) *btl-Gal4 dpp-LHG/UAS-CD8:Cherry; UAS-syt4:GFP^1-10^/lexO-CD4:GFP^11^*; (H) *btl-Gal4 dpp-LHG/+; UAS-CD4:GFP^1-10^ lexO-CD4:GFP^11^/+*; (H’) *btl-Gal4 dpp-LHG/+; UAS-CD4:GFP^1-10^ lexO-CD4:GFP^11^/UAS-syt4RNAi*; (I) *btl-Gal4 UAS-CD8GFP/+; UAS-syt1RNAi/+*; (J) *ap-Gal4/+; btl-LHG lexO-CD2:GFP/UAS-syt1RNAi*; (inset) *ap-Gal4/Gal80^ts^; dad-GFP/UAS-syt1RNAi*.

**Figure 4.** Genotypes: (A) *ap-Gal4/+; btl-LHG lexO-CD2:GFP/+*; (A’) *ap-Gal4/+; dad-GFP/+*; (B) *ap-Gal4/UAS-sybRNAi; btl-LHG lexO-CD2:GFP/+*; (B’) *ap-Gal4/UAS-sybRNAi; dad-GFP/+*; (D) *btl-Gal4 UAS-CD8GFP/UAS-sybRNAi*; *Gal80^ts^*/+; (D’) *btl-Gal4 UAS-CD8Cherry/sybRNAi; dad-GFP/Gal80^ts^*; (E) *ap-Gal4/UAS-cacRNAi; btl-LHG lexO-CD2:GFP/+*; (E’) *ap-Gal4/UAS-cacRNAi; dad-GFP/+*; (F) *ap-Gal4/UAS-stjRNAi; btl-LHG lexO-CD2:GFP/+*; (F’) *ap-Gal4/UAS-stjRNAi; dad-GFP/+*; (G) *btl-Gal4 UAS-CD8GFP/UAS-cacRNAi*; (G’) *btl-Gal4 UAS-CD8Cherry/cacRNAi; dad-GFP/+*; (H) *btl-Gal4 UAS-CD8GFP/UAS-stjRNAi*; (H’) *btl-Gal4 UAS-CD8Cherry/stjRNAi; dad-GFP/+*; (I-I’’) *UAS-CD4:GFP^1-10^ lexO-CD4:GFP^11^/+; btl-LHG bnl-Gal4/UAS-cac-TdTomato*. (J) *ap-Gal4/UAS-irk2^DN^; btl-LHG lexO-CD2GFP/Gal80^ts^*; (J’) *btl-Gal4 UAS-CD8GFP/UAS-irk2^DN^; Gal80^ts^/+*.

**Figure 5.** Genotypes: (F,G) *ap-Gal4/+; UAS-dpp/tub-Gal80^ts^*; (H-J) *btl-Gal4; UAS-GFP*;(K,L) *btl-Gal4 UAS-GCaMP6/+* for L-glutamate, and *ap-Gal4/+; btl-LHG lexO-GCaMP6/UAS-Chrimson* for photostimulation.

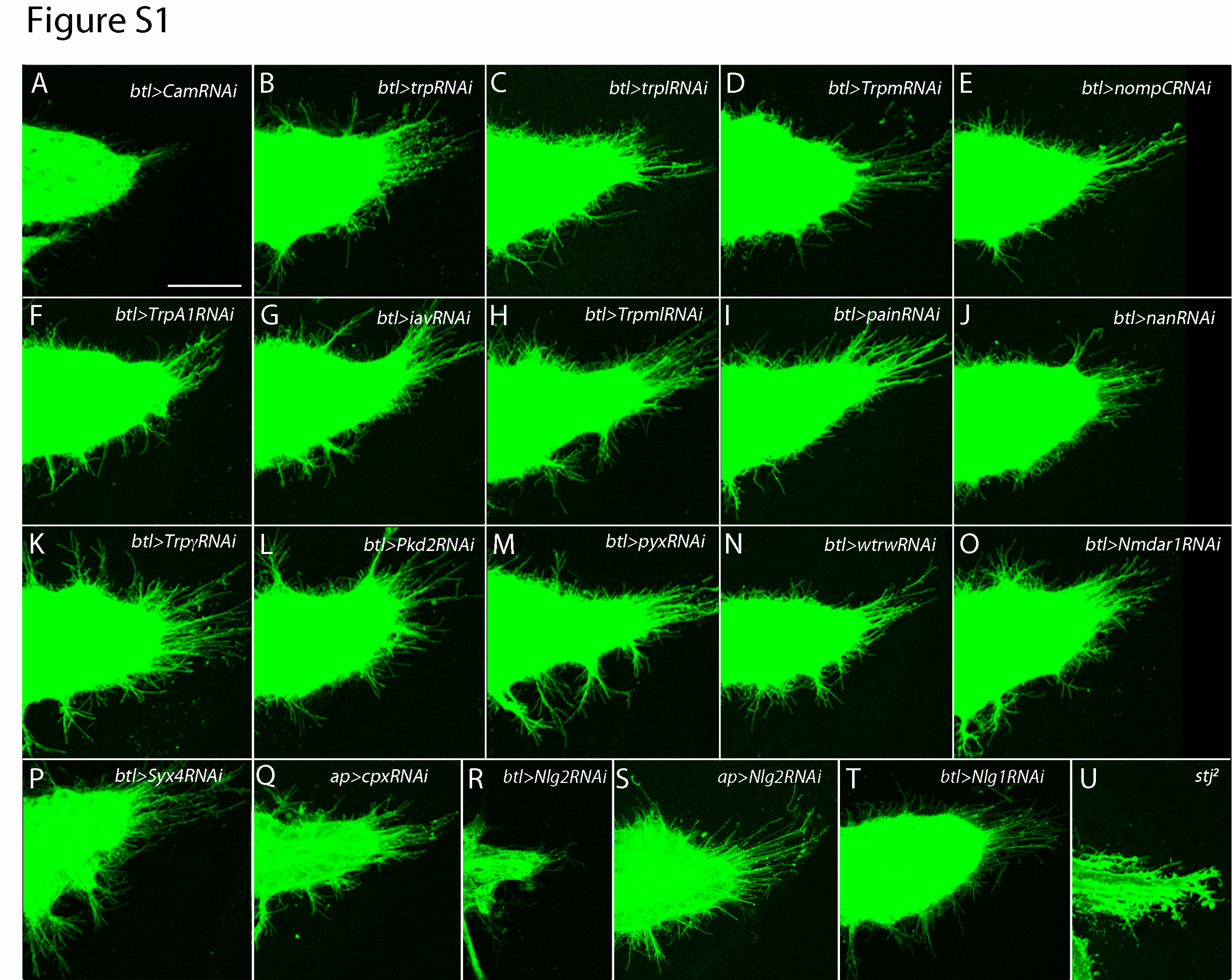

**Figure S1. ASP development in ion channel and synaptic protein mutants.** ASP morphology in animals with diminished expression of ion channels and synaptic proteins. Genotypes: (A) *btl-Gal4 UAS-CD8GFP/+ ; UAS-camRNAi/+*; (B) *btl-Gal4 UAS-CD8GFP/UAS-trpRNAi*; (C) *btl-Gal4 UAS-CD8GFP/+ ; UAS-trplRNAi/+*; (D) *btl-Gal4 UAS-CD8GFP/UAS-TrpmRNAi* ; (E) *btl-Gal4 UAS-CD8GFP/+ ; UAS-nompCRNAi/+*; (F) *btl-Gal4 UAS-CD8GFP/+ ; UAS-TrpA1/+*; (G) *btl-Gal4 UAS-CD8GFP/+ ; UAS-iavRNAi/+*; (H) *btl-Gal4 UAS-CD8GFP/+ ; UAS-trpmlRNAi/+*; (I) *btl-Gal4 UAS-CD8GFP/+ ; UAS-painRNAi/+*; (J) *btl-Gal4 UAS-CD8GFP/+ ; UAS-nanRNAi/+*; (K) *btl-Gal4 UAS-CD8GFP/+ ; UAS-trpγRNAi/+*; (L) *btl-Gal4 UAS-CD8GFP/+ ; UAS-Pkd2RNAi/+*; (M) *btl-Gal4 UAS-CD8GFP/+ ; UAS-pyxRNAi/+*; (N) *btl-Gal4 UAS-CD8GFP/+ ; UAS-wtrwRNAi/+*; (O) *btl-Gal4 UAS-CD8GFP / UAS-Nmdar1RNAi/+*; (P) *btl-Gal4 UAS-CD8GFP/UAS-Syx4RNAi/+*; (Q) *ap-Gal4/+ ; btl-LHG lexO-CD2:GFP/UAS-cpxRNAi*; (R) *btl-Gal4 UAS-CD8GFP/+ ; UAS-Nlg2RNAi/+*; (S) *ap-Gal4 UAS-CD8GFP/UAS-Nlg2RNAi*; (T) *btl-Gal4 UAS-CD8GFP/+ ; UAS-Nlg1RNAi/+*; (U) *stj^2^/stj^2^; btl-LHG lexO-CD2GFP/+*. Scale bar: 30 μm.

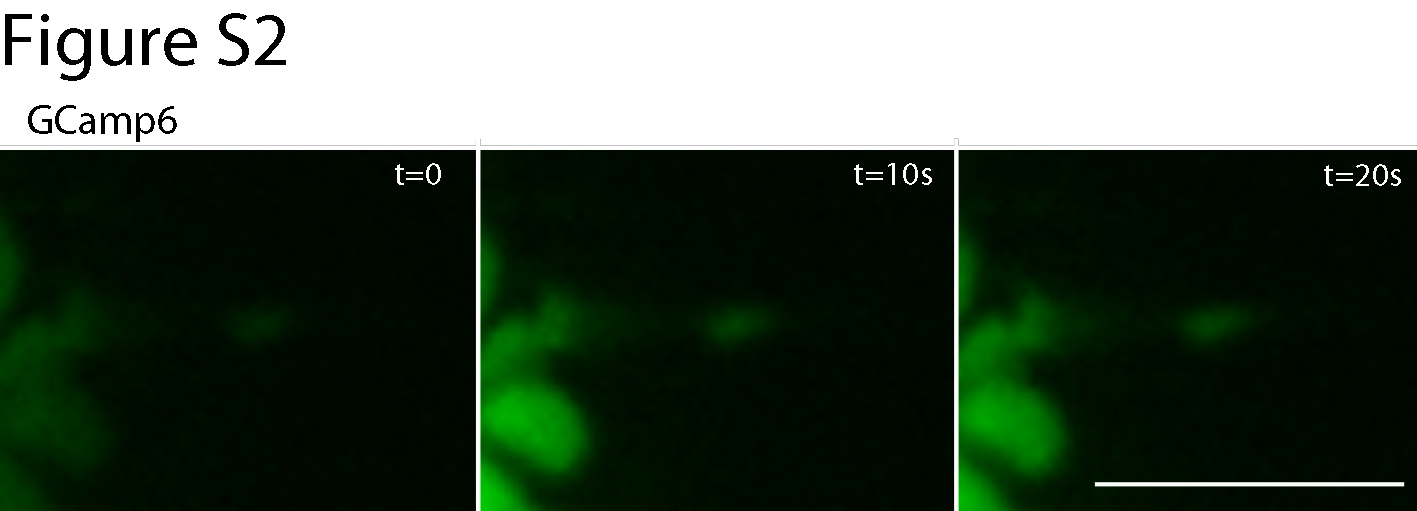

**Figure S2. Fluorescence of GCaMP6 in ASP cytonemes.** Images obtained at indicated times show GFP fluorescence detected in ASP cytonemes. Genotype: *btl-Gal4 UAS-GCaMP6/+*.

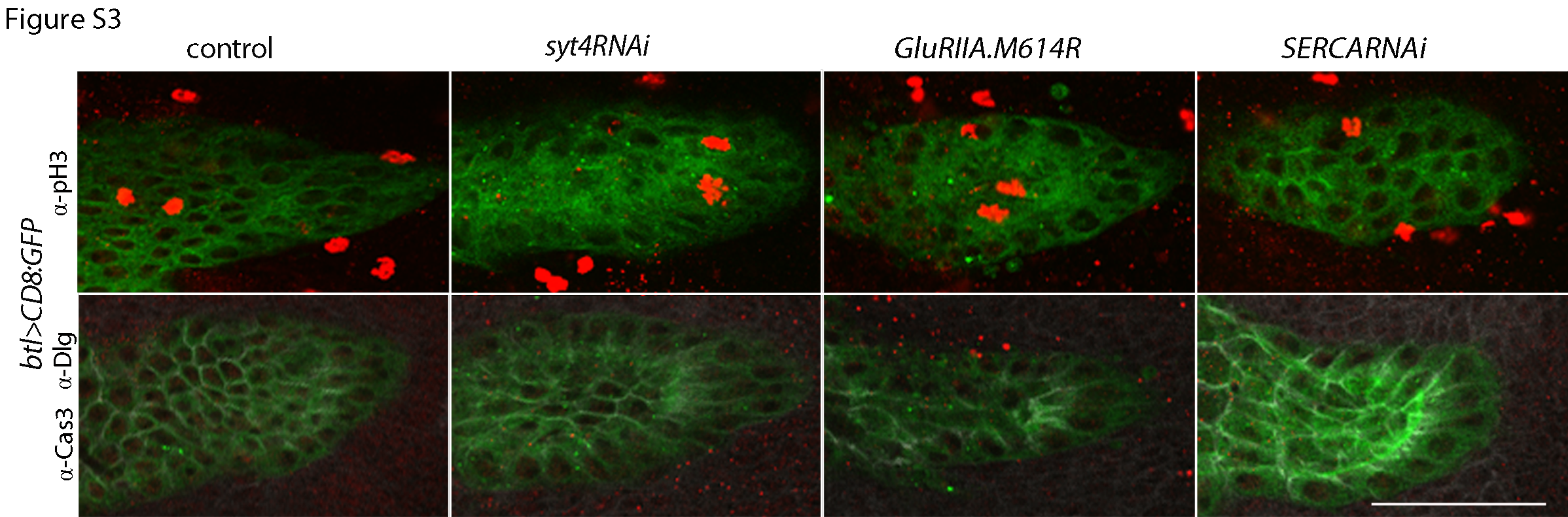

**Figure S3. Cell division and polarity in ASPs expressing *syt4RNAi*, *GluRIIA.M614R* and *SERCARNAi*.** Mitotic cells (stained with α-pH3 antibody), apoptotic cells (stained with α-Caspase-3 antibody) and cell shape (stained and α-Dlg antibody) were unchanged in ASPs downregulated for Syt4, GluRIIA or SERCA relative to control. Scale bar: 50 μm.

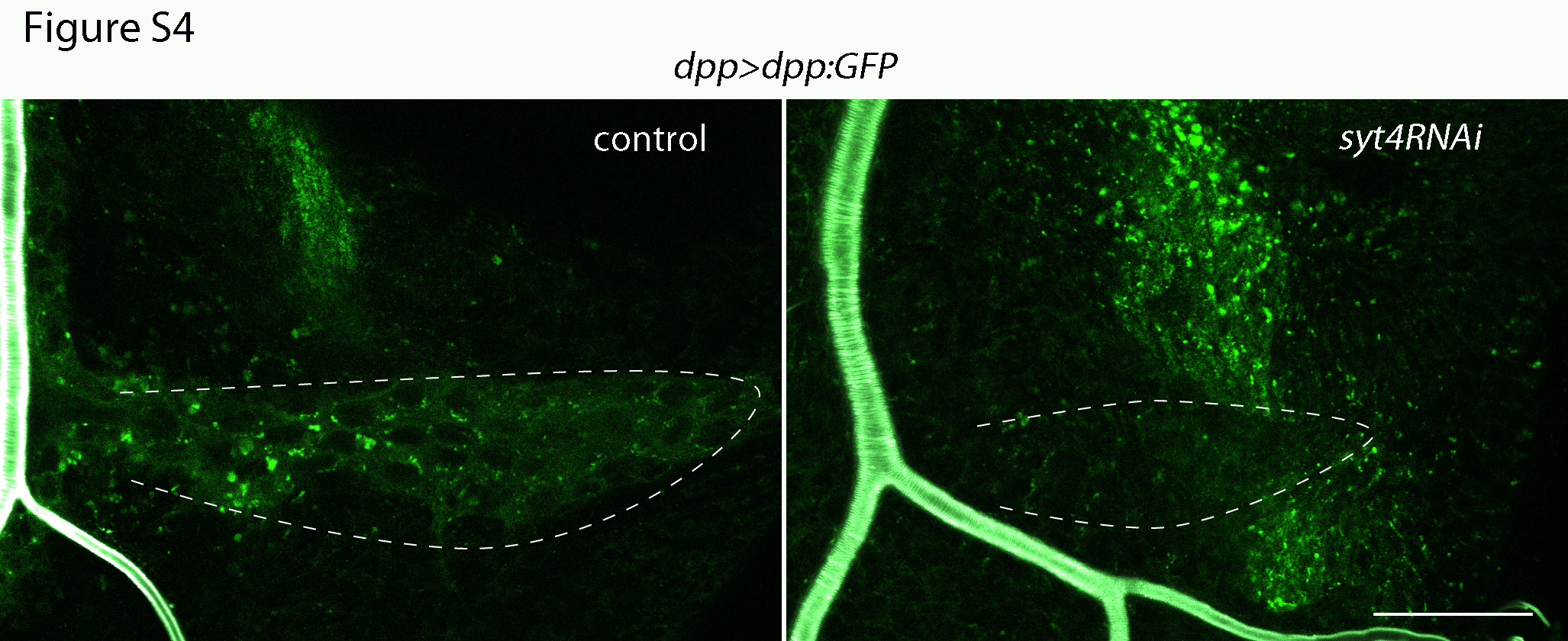

**Figure S4. Reduction of Dpp uptake in the ASP by syt4RNAi expression.** Scale bar: 30 μm.

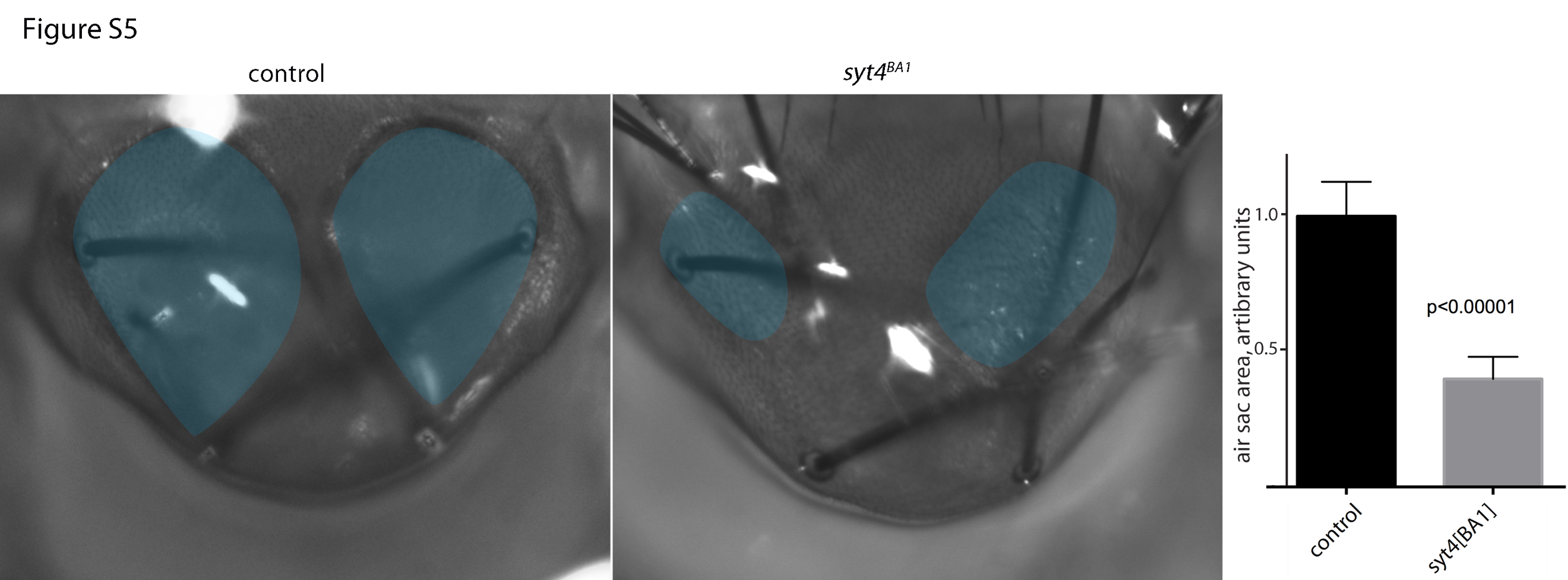

**Figure S5. Malformed dorsal air sacs in *syt4^BA1^* flies.** Images of control (left) and mutant (right) scutellums. Bar graph quantifies the relative size of control and mutant dorsal air sacs.

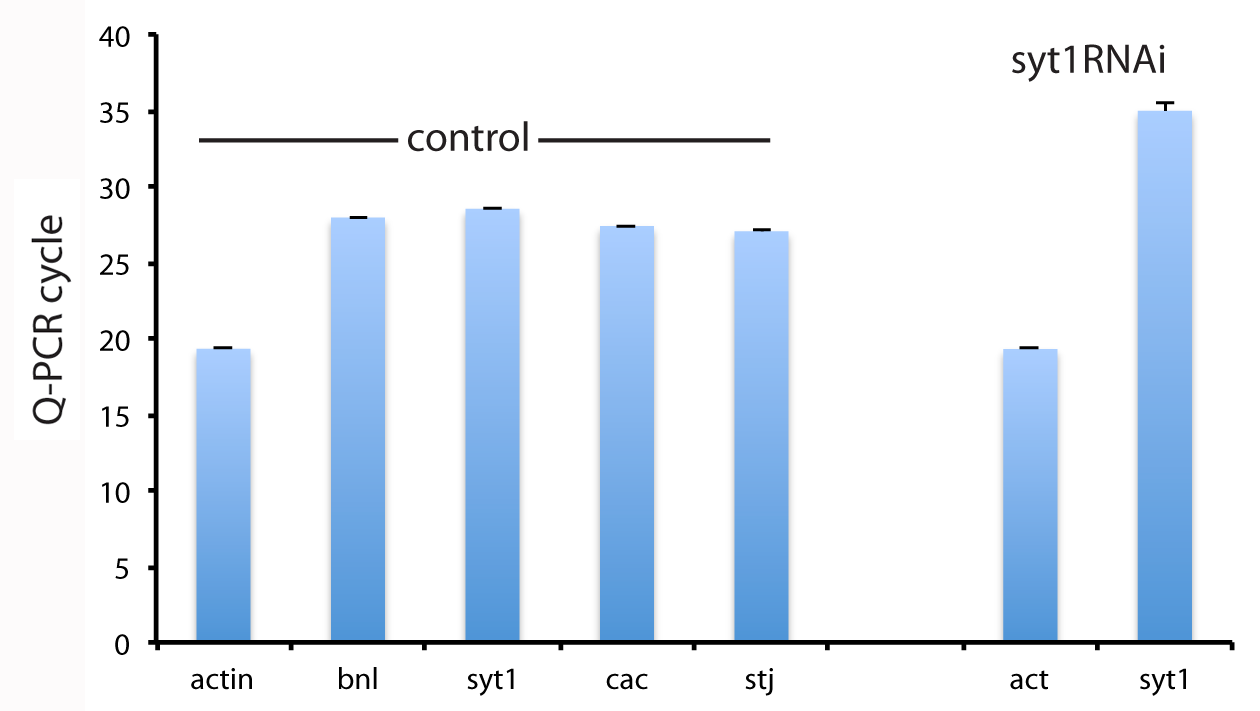

**Figure S6. Quantitative expression analysis of anterograde signaling components in the wing disc.** qRT-PCR analysis detected *syt1, cac*, and *stj* transcripts at levels approximately equivalent to *branchless*/*FGF* (*bnl*). The presence of syt1RNAi reduced the amount of syt1 RNA detected by qPCR.

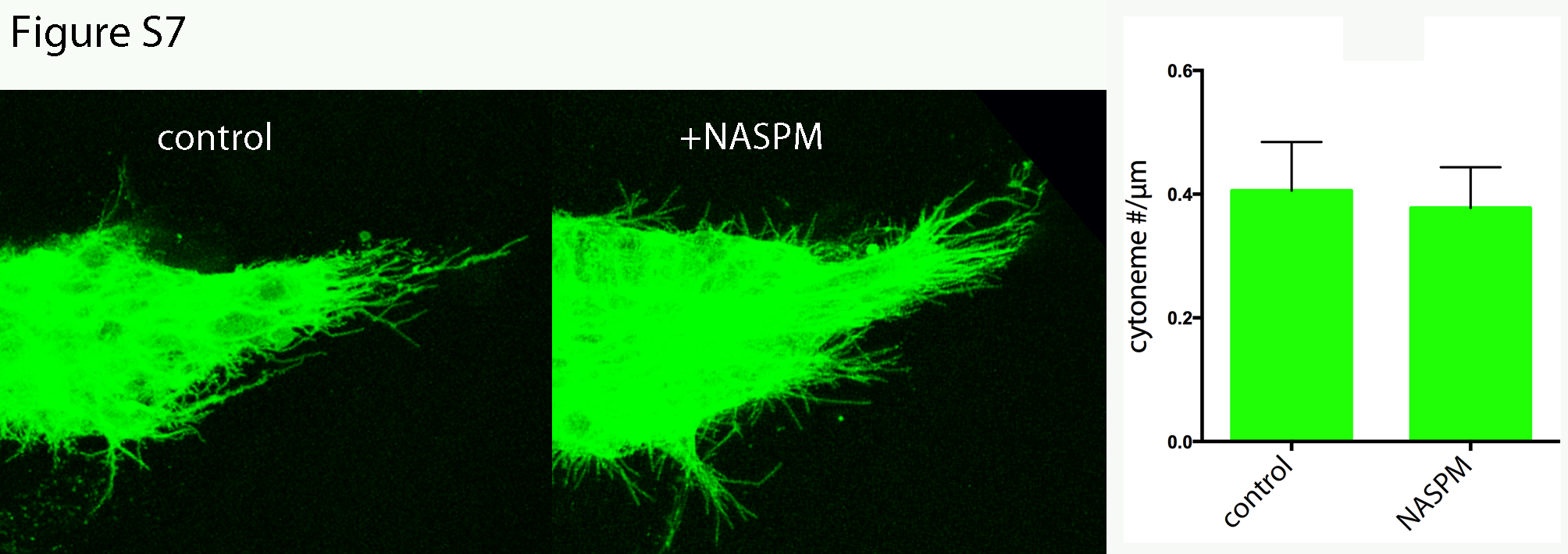

**Figure S7. Effect of NASPM on ASP cytonemes.** ASP preparations incubated for 3 hours in HL medium (left) or HL + 10 μM NASPM (right). Bar graph plots the relative number of cytonemes in control and NASPM-treated ASPs. No statistical difference was detected by Student’s t-test.

**Movie S1. Calcium transients in the ASP.** Genotype: *btl-Gal4,UAS-GCaMP6/+*

<https://drive.google.com/file/d/1197HjsGx7KOKq9XbSXbjXHFDDyTE3Xb7/view?usp=sharing>

**Movie S2. Calcium transients in ASP cytonemes containing Tkv:mCherry.** Genotype: *btl-Gal4/+; UAS-tkv:Cherry/UAS-syt4GCaMP6*.

<https://drive.google.com/file/d/1PIa285kmIz3flwkPHj080VN0yM1bI-8Z/view?usp=sharing>

**Movie S3. Calcium transients in ASP cytonemes containing Btl:mCherry.** Genotype: *btl-Gal4 UAS-btl:Cherry/+; UAS-syt4GCaMP6/+*.

<https://drive.google.com/file/d/1IsaeEVl0EcXFZdl5FnWAg6BvxIQxm2y-/view?usp=sharing>

**Movie S4. Calcium transients suppressed in ASP with downregulated SERCA** (genotype: *btl-Gal4 UAS-GCaMP6/+; UAS-SERCARNAi/+*); incubation with 2 mM EGTA, (genotype: *btl-Gal4 UAS-GCaMP6/+*); expression of glutamate receptor mutant GluRIIA.M614R, (genotype: *btl-Gal4 UAS-GCaMP6/UAS-GluRIIA.M614R*); and incubation with 10 μM NASPM.

<https://drive.google.com/file/d/1t4loyP1c9YT0u6b45PCzW2LQFOJ8I-t3/view?usp=sharing>

**Movie S5. Motile Tkv:Cherry in the cytonemes.** Genotype: *btl-Gal4 UAS-CD8GFP/+; UAS-tkv:Cherry/+*.

<https://drive.google.com/file/d/1uZUApdZGZLQmauxjTyrAW3okUVEjvGDf/view?usp=sharing>

**Movie S6. Tkv:Cherry movement reduced by presence of EGTA.** (2 mM). Genotype: *btl-Gal4 UAS-CD8GFP/+; UAS-tkv:Cherry/+*.

<https://drive.google.com/file/d/15rauPJAtnNdz_9U_bnCWQHbDXT2MziQR/view?usp=sharing>

**Movie S7. Calcium transients in the presence of 1 mM L-glutamate.** (genotype: *btl-Gal4 UAS-GCaMP6/+*).

<https://drive.google.com/file/d/1ZSXdjsd9DQVmM70D1-vyJCBiJmhxxX00/view?usp=sharing>

**Movie S8.** **Calcium transients in response to optogenetic activation.** Three minute incubation preceded a 10 sec 640nm light pulse and was followed by 8 additional minute incubation. Genotype: *ap-Gal4/+; btl-LHG lexO-GCaMP6/UAS-Chrimson*

<https://drive.google.com/file/d/1ED02TPezq7tMT4qrQAUb1Uf80DSj2yxB/view?usp=sharing>

**Movie S9**. **Motile Syt4:mRFP puncta in the cytonemes.** Syt4:mRFP expressed in trachea was detected in fluorescent puncta that move along dynamic ASP cytonemes (labeled by CD8:GFP). Genotype: *btl-Gal4,UAS-CD8:GFP/+; UAS-syt4:mRFP/+*.

<https://drive.google.com/file/d/1NipRJtAF6KiLuJna-0owRTjHC9umjt7T/view?usp=sharing>

**Movie S10. Calcium transients in the wing disc.** Genotype: *nub-Gal4/+; UAS-GCaMP6/+*.

<https://drive.google.com/file/d/1oKiCcdS-VBxaUnLCQTBiDoz-uRbmrjq0/view?usp=sharing>
